# Critical role of cell competition in gliomagenesis

**DOI:** 10.64898/2026.01.15.699808

**Authors:** Ying Jiang, Ryuhjin Ahn, Arthur Huang, Phillippe P. Gonzalez, Jungeun Kim, Guoxin Zhang, Zihao Liu, Zhenqiang He, Lindsey Dudley, Kunal S. Patel, Godfrey A. Dzhivhuho, Sam Crowl, Piotr Przanowski, Luisa Quesada Camacho, Sijie Hao, Jianhao Zeng, Simon Hippenmeyer, Mohammad Fallahi-Sichani, Kevin A. Janes, Kristen M. Naegle, Marie-Louise Hammarskjold, Steven A. Goldman, Harley I. Kornblum, Maojin Yao, Forest White, Hui Zong

## Abstract

Malignant glioma is incurable. Using a mouse genetic mosaic system to generate sporadic *Trp53,Nf1*-null OPCs, we previously identified oligodendrocyte precursor cell (OPC) as a cell-of-origin of glioma. Here, we report that pre-malignant *Trp53,Nf1*-null OPCs outcompete wildtype counterparts during their expansion. Blocking competition by mutating/strengthening wildtype OPCs impeded both pre-malignant progression and malignant expansion of glioma.

“In-tissue” phosphoproteomic profiling revealed an enrichment of phosphopeptides related to RNA splicing and protein translation at the peak of cell competition, suggesting that competitiveness may stem from unique protein species. Among candidates was mTORC1, whose pharmacological inhibition or genetic disruption resulted in a loss of competitiveness in our mouse model. Finally, analysis of patient biopsies and interrogating the role of individual gliomagenic mutations in OPC competition supported its relevance in human gliomas. Together, these findings identified the driving role of competitive interactions among OPCs in gliomagenesis, and suggest unconventional therapeutic strategies to target this process.

## INTRODUCTION

Malignant gliomas are among the most lethal human cancers. High grade gliomas – glioblastomas – are invariably fatal, while lower grade gliomas may undergo progressive anaplastic transformation to higher grade states, similarly portending death ^1,2^. The potential progression of even WHO grade 2 tumors and the post-treatment relapse despite maximal resection and aggressive treatment make glioma one of the deadliest cancer types ^1,2^. One factor underlying glioma malignancy could be the intrinsic biological robustness of its cell of origin ^3,4^. To study this problem at the pre-malignant stage of glioma, our lab previously created a mouse genetic model of glioma using the Mosaic Analysis with Double Markers (MADM) system ^5^.

Starting from a non-labeled mouse heterozygous for a tumor suppressor gene (TSG), MADM generates rare GFP-labeled TSG-null cells and RFP-labeled sibling wildtype cells through Cre/loxP-mediated mitotic recombination ^5,6^. Using MADM, we mutated *Trp53* and *Nf1*, two key TSGs involved in human gliomas ^7,8^, in all neuroglial lineages. The oligodendrocyte precursor cell (OPC) was pinpointed as the cell-of-origin in this model, based on the unique and massive expansion of GFP^+^ *Trp53,Nf1*-null OPCs over their RFP^+^ wildtype siblings at pre-malignancy ^9^. The role of OPC in glioma is broadly supported by corroborating evidence from independent studies in several labs using complementary techniques, including prevalent positive staining of OPC-specific markers in patient samples ^10–13^, prominent OPC gene expression signatures in bulk and scRNA-seq of patient samples ^4,14,15^, and through the use of many other genetic models ^16–24^. Importantly, well-studied synaptic connections between OPCs and neurons^25–29^ provided a physiological foundation for recent studies of glioma-neuron interactions ^30–33^, which have implicated neural activity as a contributor to glioma progression ^34^. Finally, studies on post-surgical tumor margins revealed the significant presence of invasive OPC-like glioma cells, making in-depth investigation of OPC biology critical for relapse prevention ^32,35,36^.

OPCs comprise the largest progenitor pool in the brain in rodents and human ^37–43^, and they remain regenerative throughout life ^44,45^. However, how pre-malignant OPCs could expand remains puzzling because normal OPCs are known to be robustly restrained by contact inhibition. Specifically, in the adult brain, OPCs form a tiling pattern, remain non-proliferative through inhibitory, transient contacts of each other’s fine processes, and only proliferate upon the death or differentiation of a neighboring OPC that locally releases the contact inhibition ^46,47^.

Furthermore, even the void-enabled proliferation is transient: OPCs promptly return to quiescence once contact inhibition is restored after cell division. Therefore, it is critical to understand the cellular mechanism that enables the pre-malignant OPCs to evade contact inhibition, which may not only explain the extraordinary malignancy of gliomas but also provide novel insights for treatment. Because the MADM model as a genetic mosaic system enabled us to study the interactions between sporadic *Trp53,Nf1*-null OPCs and normal OPCs in the same mouse brain ^5,9^, the current study reveals that, rather than passively avoiding contact inhibition, *Trp53,Nf1*-null OPCs outcompeted their wildtype counterparts during pre-malignant progression, termed OPC competition.

Cell competition is a biological process first described in epithelial tissues of *Drosophila*, defined as “survival of the *equal fitness*”: although less fit cells (potential losers) can survive among themselves as a homogenous population, they would die when they are surrounded by more fit cells (winners) in a genetic mosaic tissue ^48^. Many follow-up studies showed that uniform fitness ensured by cell competition is instrumental for both embryonic development and adult tissue homeostasis ^49–51^. However, the investigation of cell competition in mammalian cancers, especially in non-epithelial tissues, remains at its infancy because of the scarcity of mammalian genetic mosaic models available for studying the competitive process, as well as the technical challenge of pinpointing competition-specific signaling mechanisms without disrupting cell-cell contacts.

To determine the functional relevance of our discovery in the context of glioma, we blocked the competitiveness of *Trp53,Nf1*-null OPCs by introducing a mutation to strengthen wildtype OPCs that results in their survival. The neutralization of cell competition not only prevented pre-malignant progression, but also impeded the malignant expansion of glioma cells, demonstrating its critical role of OPC competition in glioma. To uncover signaling mechanisms that enable mutant OPCs to outcompete their wildtype counterparts, we used an “in-tissue” phosphoproteomic profiling platform ^52^ to avoid the disruption of cell-cell interactions during tissue dissociation. After filtering out noise from other brain cells with age-matched normal brains, we focused on temporal patterns of signaling events that match the peak age of OPC competition, and discovered that competition-specific signaling nodes appear to generate unique protein species through the regulation of mRNA splicing and protein translation. Importantly, one of the candidates, mTORC1, was successfully validated by both *ex vivo* and *in vivo* assays for its specific role in OPC competition but not in other aspects of OPC biology. Finally, we established the likely importance of OPC competition in human gliomagenesis by reporting a paucity of OPCs in patient biopsies, and by individually testing a broad range of patient-related gliomagenic mutations for their contribution to OPC competition. Taken together, this study reveals OPC competition as a driving force of malignant progression of OPC-related gliomas, and sheds light on paradigm-shifting therapeutic concepts to tame glioma malignancy through neutralizing OPC competition.

## RESULTS

### *Trp53,Nf1*-null (TN-null) OPCs outcompeted normal OPCs at pre-malignancy

To investigate how pre-malignant OPCs overcome contact inhibition for their expansion, we used an OPC-specific NG2-Cre in the MADM-*Trp53,Nf1* (MADM-TN) model to generate sporadic GFP^+^ *Trp53,Nf1*-null OPCs in the brain, herein referred to as “TN-null OPCs” (Figure 1A, see S1 for detailed scheme of MADM). For the sake of simplicity, we referred to the remaining OPCs in the brain (sporadic RFP^+^ *Trp53,Nf1*-wildtype OPCs, sporadic GFP^+^RFP^+^ *Trp53,Nf1*-heterozygous OPCs, and colorless *Trp53,Nf1*-heterozygous OPCs) as “normal OPCs” hereafter. A time course analysis from postnatal day 10 to 60 (P10 to P60) showed that initially sporadic GFP^+^ TN-null OPCs expanded gradually until they populated the entire brain (Figure 1B, C). Correspondingly, the ratio between GFP^+^ TN-null and sibling RFP^+^ wildtype OPCs (G/R ratio) increased from the initial 1:1 to nearly 300-fold at P60 (Figure 1E, top). Then we realized that, if TN-null OPCs simply ignored the contact inhibition to reach such a high G/R ratio, the MADM brains would have been filled with an enormous number of OPCs. However, our staining for OPC distribution in the brain contradicted this prediction since the overall OPC density only increased marginally throughout pre-malignant expansion of TN-null OPCs (Figure 1D). Quantification further confirmed that the overall OPC density merely increased by ∼2-fold at P60 (Figure 1E, bottom), dramatically lower than the ∼300-fold increase of the G/R ratio) (Figure 1E, top). While our data strongly suggests that TN-null OPCs likely outcompeted normal OPCs during their expansion, we could not completely exclude the alternative interpretation that TN-null OPCs simply hindered the expansion of normal OPCs when the brain increases its size during postnatal development. To definitively rule out this alternative interpretation, we quantified the absolute number of normal OPCs in the brains at various ages, predicting a gradual decline if the normal OPCs were eliminated as a result of cell competition. This was exactly what we observed after systematically quantifying the absolute number of normal OPCs in ≥ 4 brains at each indicated age (Figure 1F). Next, we used immunohistological analysis to determine if TN-null OPCs specifically outcompete normal OPCs or indiscriminately harm all brain cell types, and found that only normal OPCs were uniquely absent (Figure 1G) but other brain cell types showed no detectable changes while intermixing with TN-null OPCs (Figure S2). Taken together, our finding matches the definition of cell competition ^48^: <u>winner cells specifically outcompete loser siblings</u> while coexisting in the same tissue, although the latter can survive just fine in the absence of the former. Because cell competition in this study occurs specifically between TN-null and normal OPCs, we referred to it as OPC competition hereafter.

**Figure 1.**
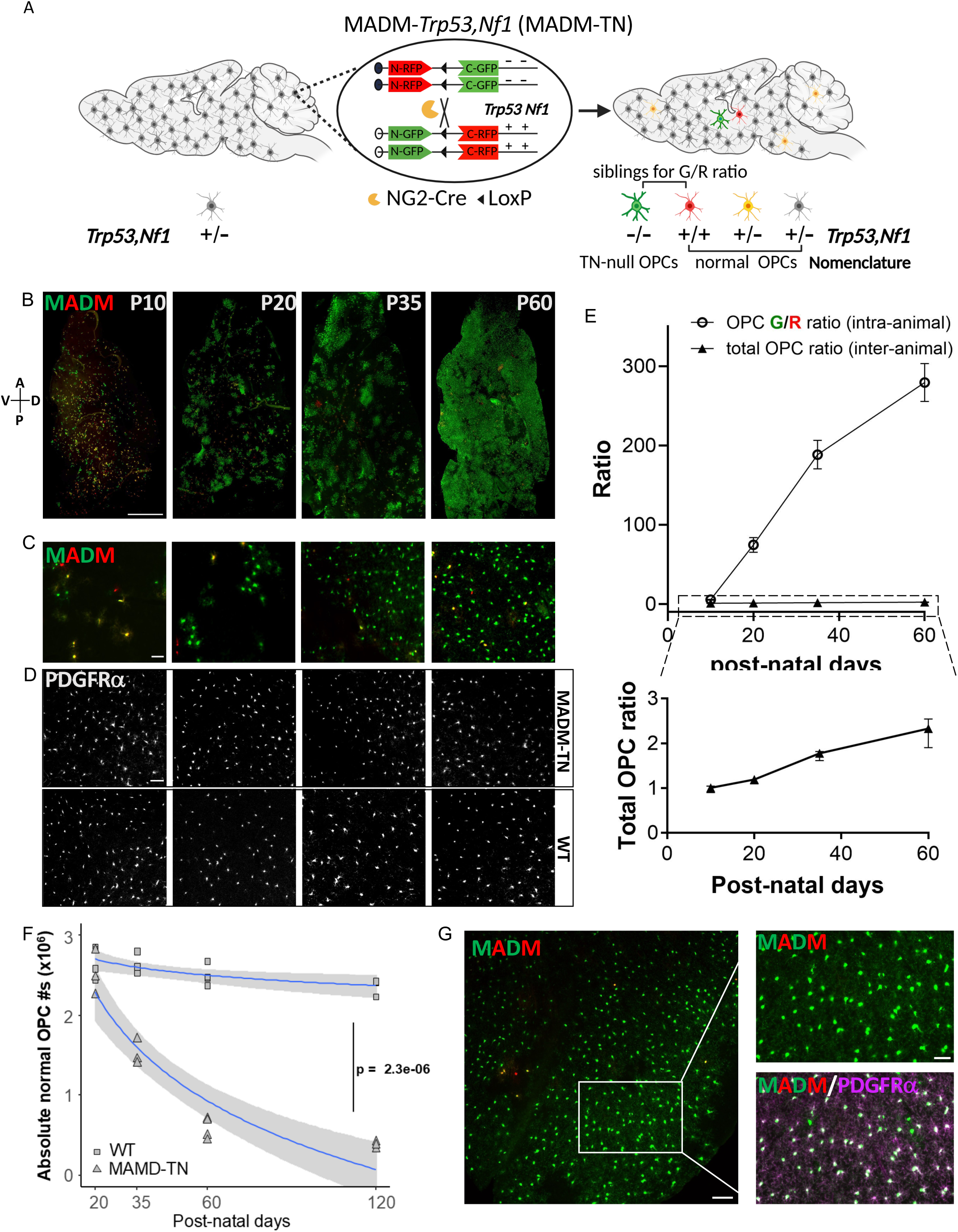
*Trp53,Nf1*-null OPCs outcompete their neighboring normal OPCs. **A.** Sporadic generation of GFP^+^ *Trp53,Nf1*-null and RFP^+^ wildtype OPCs in the MADM-TN mouse model. Ratio of sibling pair (G/R ratio) assesses the expansion of *Trp53,Nf1*-OPCs. *Trp53,Nf1*-null OPCs are herein referred to as “TN-null OPCs”; and OPCs of all other genotypes as “normal OPCs”. **B.** Macroscopic images of entire sagittal sections of MADM-*Trp53,Nf1* (MADM-TN) brains at various ages, showing the gradual expansion of TN-null OPCs. Scale bar: 1000 μm. **C.** High-magnification images showing MADM labeled OPCs at various ages. Scale bar: 50 μm. **D.** High-magnification images showing the total OPCs (PDGFR⍺^+^) in MADM-TN and age-matched wildtype brains. Scale bar: 50 μm. **E.** Line graph showing >300-fold increase of G/R ratio from P10 to P60 in MADM-TN brains (≥5 brains per age group) but only ∼2-fold increase of total OPC number in MADM-TN brains as compared to age-matched wildtype brains during the same period (≥3 brains per age group). **F.** The absolute number of normal OPCs declines dramatically with time in MADM-TN mice. Temporal trends in normal OPC numbers from n ≥ 4 MADM-TN and wildtype animals per time point were modeled as a genotype-dependent exponential decay. The p value reports the statistical significance of the genotype term in the log-transformed linear model. **G.** GFP^+^ TN-null OPCs out-compete and replace normal OPCs at P60. Representative image on the left showing the presence of a large number of TN-null OPCs. Scale bar: 100 μm. High-magnification images on the right showing a nearly complete absence of wildtype OPCs. Scale bar: 50 μm.

Since the MADM model generated TN-null OPCs at the prenatal/neonatal age, there is a possibility that OPC competition only occurs during brain development. If this were the case, OPC competition may not be relevant in human gliomas, which mostly occur in older patients. To determine if OPC competition also occurs in the adult brain, we generated sporadic *Trp53,Nf1*-null OPCs in adult mice with a *NG2-CreER; Trp53flox,Nf1flox; ROSA26-LSL-tdTomato* mouse model, herein referred to as CKO (Figure S3A)^53^. At P21, a low dosage of tamoxifen was given to both control (*NG2-CreER; ROSA26-LSL-tdTomato*) and CKO mice that resulted in a small number of tdTomato^+^ wildtype and TN-null OPCs, respectively. A time course analysis was performed at 2-days post-injection (dpi), 60 dpi, 120 dpi, and 180 dpi (Figure S3B), showing that, while tdT^+^ wildtype OPCs failed to expand in the control group, tdT^+^ TN-null OPCs greatly expanded their numbers as the mice aged (Figure S3C). To determine if the expansion of TN-null OPCs was a result of cell competition, we systematically quantified unlabeled, wildtype OPCs in the CKO model across ages. The gradual decline of wildtype OPCs (Figure S3D), similar to what was observed in MADM-TN model (Figure 1G), suggests that pre-malignant OPC-normal OPC competition is not limited in the milieu of brain development, but rather occurs in an ongoing fashion the adult brain during homeostasis.

### Comprehensive analysis of the process of OPC competition

To understand the kinetics of OPC competition, we comprehensively characterized the expansion of TN-null OPCs at different ages in the MADM-TN model. At P10, TN-null OPCs almost always existed as GFP^+^ singlets; by P20, the start of OPC competition was marked by small, discrete GFP^+^ clusters, likely representing individual clones rather than aggregations of independently generated TN-null OPCs since they are rare; by P35, GFP^+^ clones were sizable but still mostly discrete; and by P60, GFP^+^ clones merged with each other to occupy almost the entire brain (Figure 2A). Such clonal expansion was evident across all brain regions, including both gray and white matter (Figure S4, representative images at P21). The range of OPC competition appeared to be short because we did not observe disrupted pattern of normal OPCs intermixing with TN-null OPCs, which would result from long-range competition; instead, we observed a nearly total exclusion of normal OPCs in GFP^+^ clones that implies local competition at or near clonal boundary (Figure 2B). We next wondered if TN-null OPCs are relatively equal (i.e. all clones can expand) or have intrinsic hierarchies (i.e. only some clones can expand) (Figure 2C). The answer to this question required the assessment of individual clonal sizes at various ages, for which we developed a workflow that involves tissue clearing, light-sheet 3D imaging, and computational analysis (Figure S5). To ensure the accuracy of measurement of clonal sizes, we focused our analysis on the cortical regions where clones are mostly isolated from each other (Supplemental Video). Using the Density-based Spatial Clustering of Applications with Noise (DBSCAN) algorithm (see method), our analysis showed that, while clones varied in size even at the same age, likely because MADM-based recombination occurred asynchronously, they collectively trended larger with age (Figure 2D), suggesting that all clones can expand. Finally, to determine if the kinetics of clonal expansion depend on clonal size (i.e. larger clones may exert stronger collective power to compete), we employed a mathematical modeling method (see details in methods) (Figure 2E). Based on the clonal distribution of P15 brains, we tested two models of expansion that predict the size distributions at P21 and P28. We found that the model with the additional assumption that growth rate depends on clonal sizes failed to outperform the simpler model that only assumed a variable growth rate independent of clone size. Taken together, our data demonstrated that all TN-null OPC clones can compete and expand, and that the rate of expansion is not dependent on clone size, suggesting that OPC competition is a prevalent and steady process.

**Figure 2.**
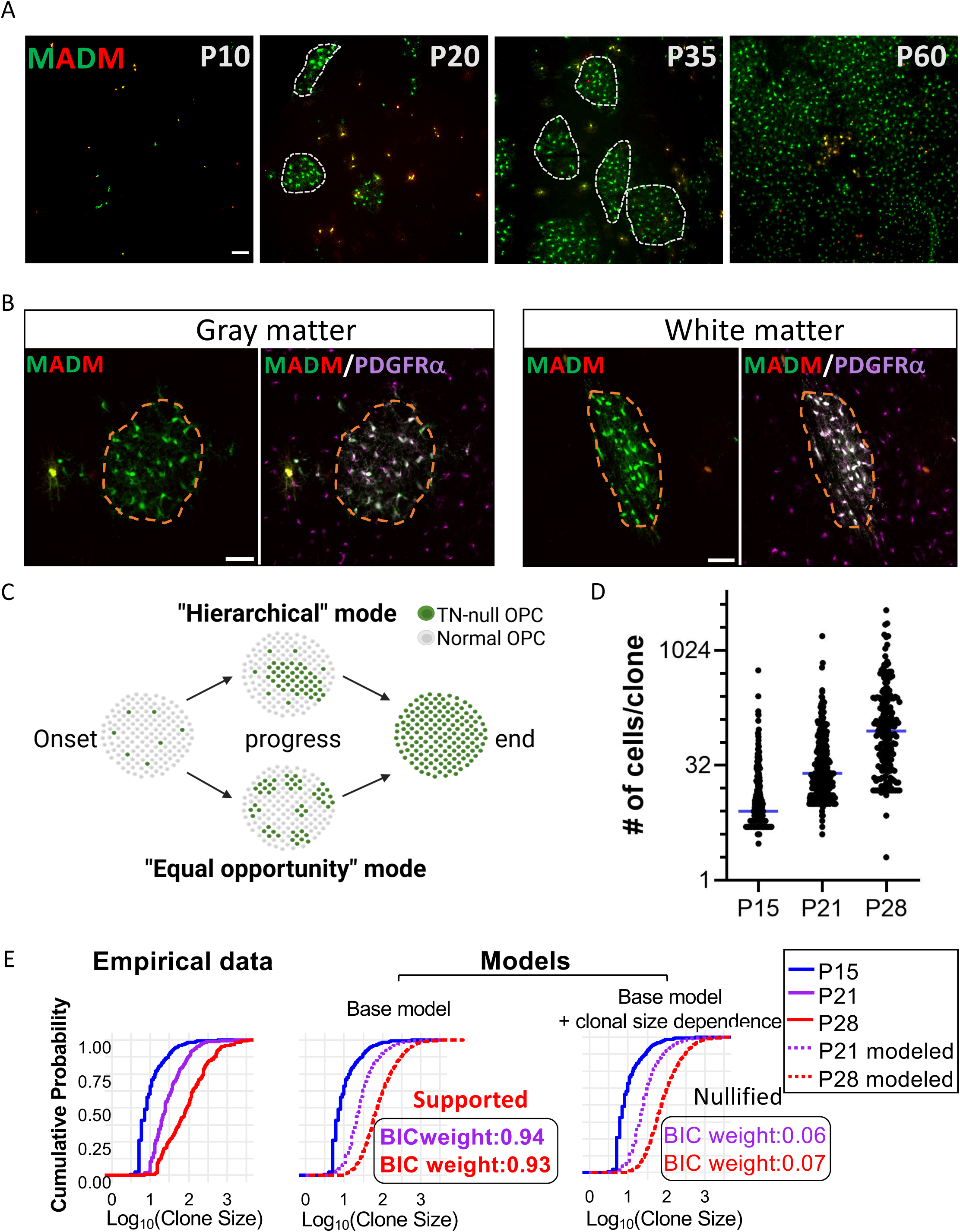
Analysis and modeling of the clonal expansion of TN-null OPCs during competition. **A.** Representative images of TN-null OPC clones in the cortical region of MADM-TN brains at various ages. White-dotted lines outline the boundary of the clones. Scale bar: 100 μm. **B.** Representative images showing the complete absence of normal OPCs in GFP^+^ TN-null OPC clones at both gray and white matters of MADM-TN brains. Scale bar: 50 μm. **C.** Schematic of two modes of TN-null OPC clonal expansion. **D.** Clonal size (3-D) distribution in the cortical regions of P15, P21 and P28 MADM-TN brains, **E.** Clonal size distributions are captured by a minimal model of variable growth rate without positive reinforcement by colony size. Clones were simulated stochastically using a baseline growth rate with a Poisson-distributed noise term. The alternative model further increases the growth rate log-linearly with colony size. The two models were compared by Bayes Information Criterion (BIC) weights (see Methods).

### Blocking OPC competition not only prevented pre-malignant progression, but also markedly impeded malignant expansion of glioma

A key question about our discovery is whether or not the competition between pre-malignant TN-null and wildtype OPCs is critical for gliomagenesis. To answer this question, we first teased apart the contribution of two gliomagenic mutations to OPC competition by establishing new MADM models with single *Trp53*-mutant (MADM-*Trp53*) and single *Nf1*-mutant (MADM-*Nf1*), respectively (Figure S6). MADM-*Trp53* phenocopied MADM-WT brains while MADM-*Nf1* model phenocopied MADM-TN (Figure 3A, B), suggesting that the loss of *Nf1* is solely responsible for OPC competition in our model. Interestingly, among twelve MADM-*Nf1* mice that we examined at or after the latency age, none formed malignant gliomas, indicating that *Nf1*-loss mediated OPC competition is not sufficient for malignant transformation (Figure 3C, D). The discovery of *Nf1*-null OPCs as “competitive but not transformative” provided an opportunity for us to determine whether blocking OPC competition by pitting *Nf1*-null OPCs against *p53,Nf1*-null OPCs could prevent gliomagenesis. To test whether *Nf1*-null OPCs can resist the competition of TN-null OPCs, thereby restraining the expansion of TN-null OPCs and eventual malignant transformation, we established a MADM-anti-competition (MADM-AC) model. The MADM-AC model differs from the original MADM-TN model only by recombining an *Nf1*-flox allele onto the previously wildtype MADM allele (Figure 3F vs. E, Figure S7). In this model, OPC-specific NG2-Cre catalyzes MADM recombination to generate rare, GFP^+^ *Trp53*,*Nf1*-null OPCs in the brain, while also catalyzing conventional conditional knockout to turn *Nf1*-flox/*Nf1*-flox into *Nf1*-null in all OPCs. Intriguingly, prevalent *Nf1*-null OPCs neutralized the competitive advantage of TN-null OPCs, resulting in a G/R ratio of ∼1 (Figure 3B) and a complete lack of clonal expansion of GFP^+^ TN-null OPCs (Figure 3H vs. G). Finally, we examined ten MADM-AC mice at P240 (tumor latency age of this model) and found that none of them had gliomas (Figure 3D, I-K). This proof-of-principle experiment clearly demonstrated that OPC competition is a necessary antecedent to pre-malignant expansion and ensuing malignant transformation.

**Figure 3.**
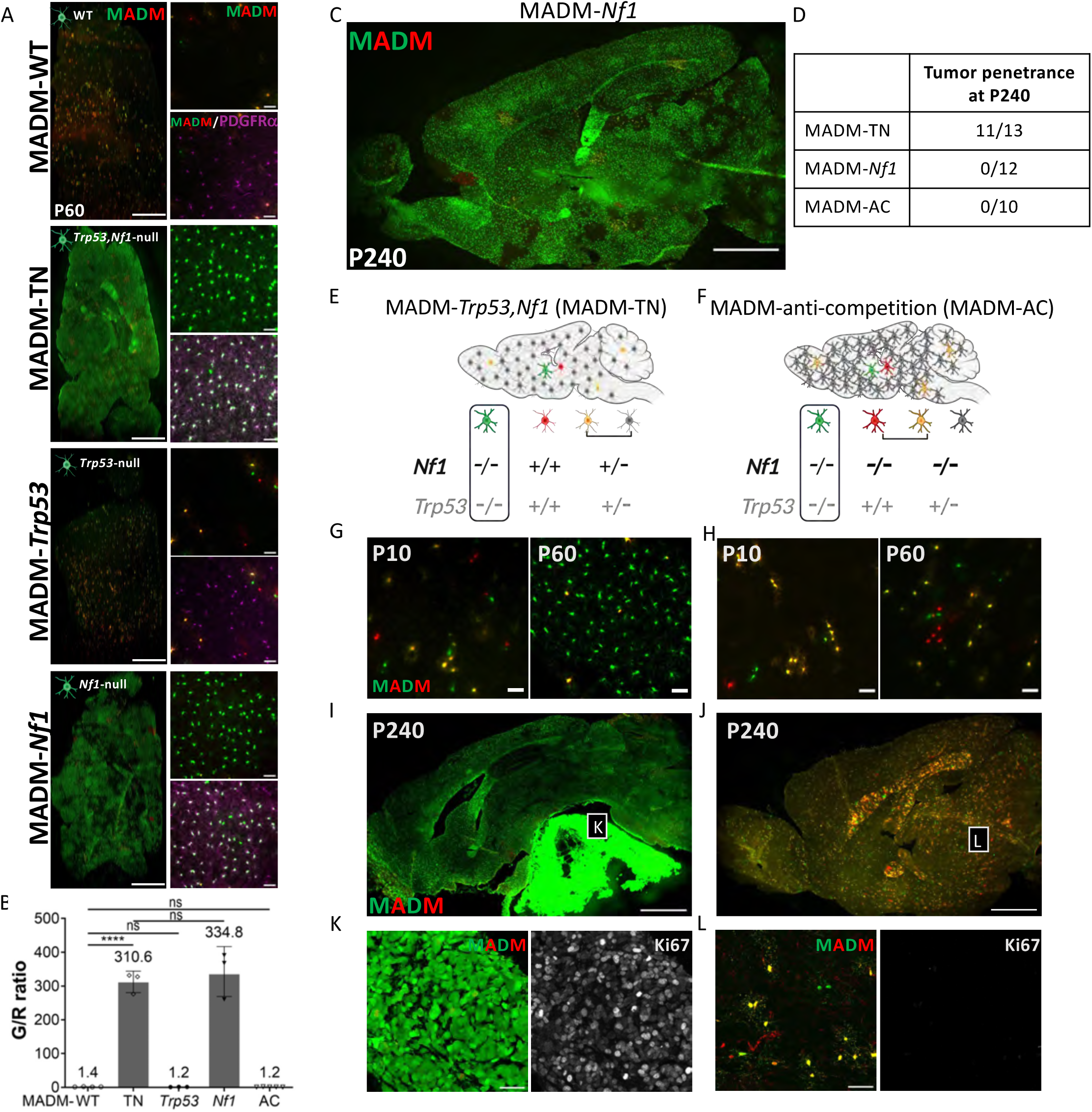
Blocking OPC competition prevents the pre-malignant expansion of TN-null OPCs. **A.** Macroscopic images of sagittally sectioned P60 brains from MADM-WT; MADM-TN; MADM-*Trp53*; MADM-*Nf1* mice (left, scale bar: 1000 μm); high-magnification images on the right showing the competition or lack of competition by mutant OPCs, scale bar: 50 μm. **B.** Bar graph showing the G/R ratio of OPCs in MADM-WT; MADM-TN; MADM-*Trp53*; MADM-*Nf1*; and MADM-AC models. Ratio t-test (using geometric mean) was performed to compare the G/R ratio between different models. ****p<0.0001, ns p>0.05. **C.** Macroscopic image of sagittally sectioned brains from MADM-*Nf1* mice at P240, showing that *Nf1*-null OPCs fully occupied the brain parenchyma but did not transform malignantly. Scale bar: 1000 μm. **D.** Tumor penetrance in the MADM-TN, MADM-*Nf1*, MADM-AC models. **E,F.** Genotypes of OPCs in MADM-TN and MADM-anti-competition (AC) brains, respectively. **G,H.** Representative images of expansion of TN-null OPC from P10 to P60 in the MADM-TN brains (**G**), and the lack of expansion in MADM-AC brains (**H**), respectively. Scale bar: 50 μm. **I-L.** Macroscopic image of sagittal section of MADM-TN brains at P240 (**I**), revealing a GFP^+^ proliferating (Ki67^+^) glioma (**K**). Representative macroscopic image of sagittal section of MADM-AC brains at P240 (**J**), showing the rarity of GFP^+^ and Ki67^+^ cells (**L**). Scale bar in **I,J**: 1000 µm; scale bar in **K,L**: 50 μm. These are representative images of ≥ 10 brains from MADM-TN and MADM-AC mice, respectively.

Since our data above solely focused on pre-malignancy, next we wondered whether transformed glioma cells still compete with normal OPCs, and if so, whether blocking OPC competition could hinder malignant progression. To address the first question, we performed a syngeneic grafting experiment with glioma cells isolated from the MADM-TN model (Figure 4A). We assessed the abundance of host OPCs in the tumor region (Figure 4B), focusing on tumor boundaries where glioma cells interact with host cells in the brain parenchyma. In comparison to the contralateral normal brain region, the number of OPCs dropped significantly in the tumor region (Figure 4C-E) while neurons and astrocytes remained similar (Figure 4F-H, I-K), suggesting that grafted glioma cells specifically out-competed host OPCs when they infiltrated into the brain parenchyma. To determine if *Nf1*-null OPCs can restrain malignant glioma cells, we grafted glioma cells into host brains containing wildtype OPCs versus *Nf1*-null OPCs, and performed analysis at 5 weeks post grafting (Figure 4L). Strikingly, glioma sizes were markedly smaller in host brains containing *Nf1*-null OPCs compared to those grafted in host brains containing wildtype OPCs (Figure 4M,N). To determine how *Nf1*-null OPCs restrain glioma cells, we first examined the proliferative rates of infiltrative glioma cells grafted in two different host brains and found they remained similar (Figure 4O,P), suggesting that the restraint of glioma expansion is unlikely intrinsic to glioma cells. In contrast, the dispersion of grafted glioma cells in brains with *Nf1*-null OPCs, when compared to those grafted in wildtype host brains, was greatly impeded (Figure 4Q,R). Taken together, these findings suggest that the subdued expansion of glioma in the host brains with *Nf1*-null OPCs was caused by the restriction of glioma infiltration by competitive *Nf1*-null OPCs, rather than caused by the decrease of intrinsic proliferative potential in glioma cells, supporting the notion that blocking OPC competition can hinder malignant progression.

**Figure 4.**
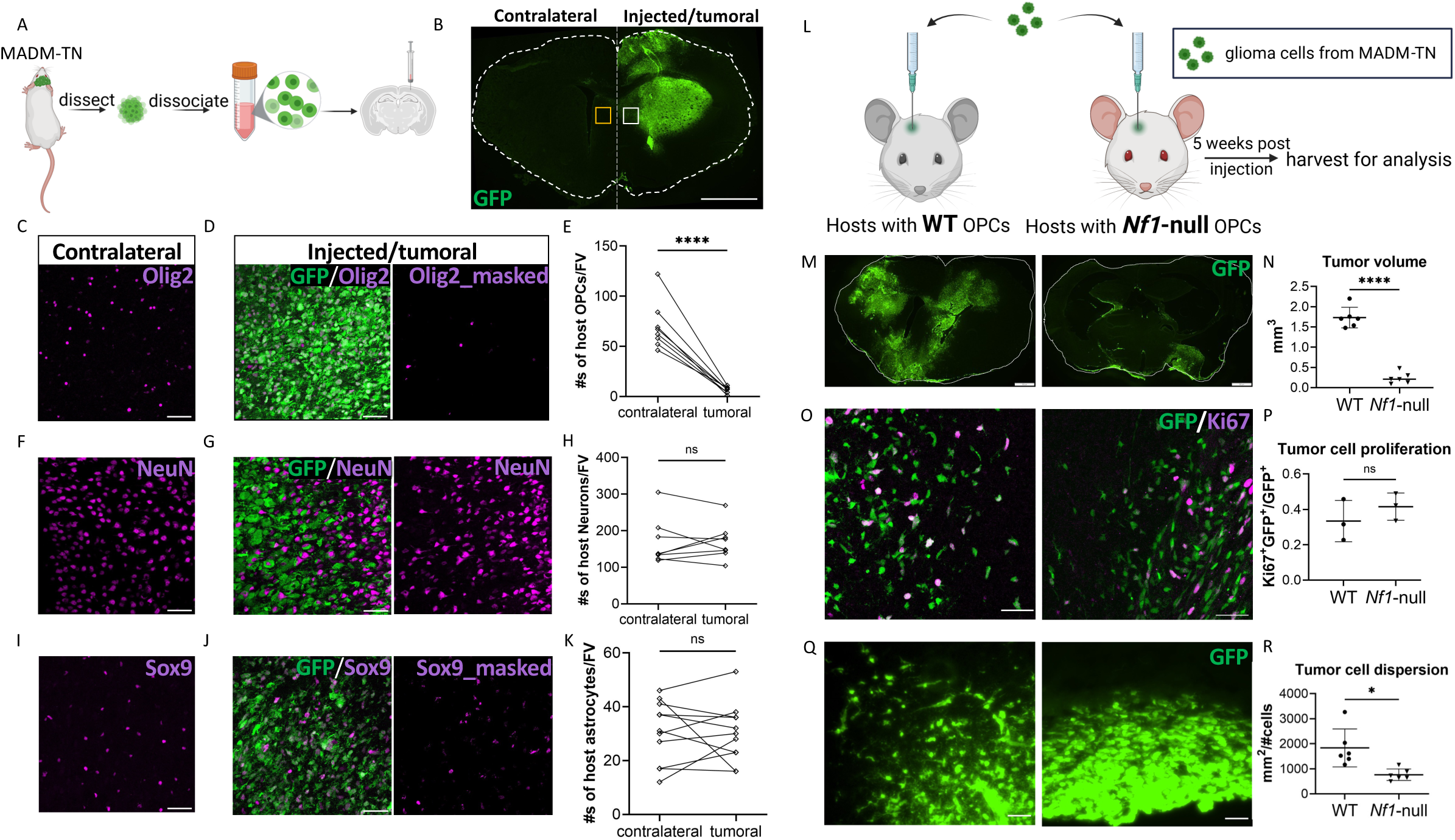
Blocking OPC competition markedly impedes the expansion of malignant glioma cells. **A.** Workflow for the isolation of mouse glioma cells and orthotopic grafting into mouse brains. **B.** Macroscopic image of coronally sectioned mouse brain showing the tumor grafted region (GFP^+^, right side) and tumor-free contralateral side (left side). Scale bar: 1000 µm. **C-K.** Representative images showing host OPCs (Olig2^+^) in the contralateral side (**C**) and tumor grafted side (**D**); host neurons (NeuN^+^) in the contralateral side (**F**) and tumor grafted side (**G**); host astrocytes (Sox9^+^) in the contralateral side (**I**) and tumor grafted side (**J**). Due to mouse glioma cells originating from OPCs, GFP^+^Olig2^+^ tumor cells were “masked” in panel **D** for a clearer view of host OPCs in the tumor regions. Similarly, we found that mouse glioma cells also express Sox9 and thus GFP^+^Sox9^+^ tumor cells were “masked” in panel **J** to show host astrocytes. At least 8 paired regions from 4 grafted mice were sampled for the quantification (**E,H,K**). Pair-wise t-test was performed. **** p<0.0001, ns p>0.05. Scale bar: 50 μm. **L.** Workflow of orthotopic grafting of isolated mouse glioma cells into host brains containing either WT or *Nf1*-null OPCs (n = 6 in each group). **M.** Representative images showing the growth patterns of grafted glioma cells in host brains. Scale bar: 1000 µm. **N.** The tumor volume was measured by quantifying GFP^+^ area across sections of ∼300 µm. ****p<0.0001, t-test was performed. **O.** Representative images showing the proliferation (Ki67^+^) of grafted glioma cells at the tumor boundary. Scale bar: 50 μm. **P.** 5-6 fields of grafted glioma cells in the brain parenchyma per brain were sampled. Proliferation rate of grafted glioma cells was calculated by dividing the sum of proliferative (Ki67^+^GFP^+^) cells with the total number of glioma cells (GFP^+^). 3 mice per group were used for quantification. ns p>0.05, t-test was performed. **Q.** Representative images showing the dispersion of glioma cells at the tumor boundary. Scale bar: 50 μm. **R.** Tumor cell dispersion was measured by dividing unit area with the number of GFP^+^ tumor cells within 200 µm from the boundary. *p<0.05, t-test was performed.

### Aberrantly activated Ras signaling in TN-null OPCs conferred competitiveness against wildtype OPCs

Next, because Nf1 has many functional domains beyond the well-known Ras-GAP domain that inactivates Ras (Figure S8A), we asked whether the over-activation of Ras or other dysregulated biological processes caused by the loss of *Nf1* were responsible for OPC competition. To answer this question, we established a MADM-*Nf1*^GRD^ model, in which a R1276P point mutation was made in the GAP related domain (GRD) of Nf1 to prevent it from inactivating Ras (Figure S8B, C). The fact that this model phenocopied MADM-TN and MADM-*Nf1* (Figure S8D) suggests that over-activated Ras due to the loss of GAP activity of Nf1 is responsible for OPC competition.

Since the aberrant activation of Ras leads to OPC competition, we set out to establish an *ex vivo* platform to test the ability of Ras pathway inhibitors to weaken the competitiveness of TN-null OPCs. Because the motility of cells in a dish limits the contact between TN-null and WT OPCs and thus reduces the robustness of the assay, we established an organotypic hippocampal slice platform, in which the contact between TN-null and WT OPCs are maintained within the slice ^54^. First, we validated clonal expansion in the long-term cultured MADM-TN hippocampal slices (Figure S9A, top row). Next, we tested whether MEK inhibition could weaken the competitiveness of TN-null OPCs because MEK is a key downstream effector of Ras and MEK inhibitors (MEKis) can be used to treat Ras-related cancers ^55^. Among MEKi’s, we chose Trametinib because it is a well-studied small-molecule inhibitor ^56^ that has been approved for use in several indications in human cancers alone ^57,58^ or in combination with various other inhibitors ^57,59–62^. We observed that, at a low dosage (10nM), Trametinib failed to block the expansion of TN-null OPCs (Figure S9A, middle row, and B); at a higher dosage (50nM), the apparent effect of Trametinib to block the expansion of TN-null OPCs (Figure S9A, bottom row, and B) was caused by its general toxicity to all OPCs, resulting in its dramatically decreased OPC density (Figure S9C, bottom). The fact that MEKi Trametinib failed to specifically block OPC competition motivated us to consider phosphoproteomic profiling to identify other Ras-mediated, competition-specific signaling pathways.

### “In-tissue” phosphoproteomic profiling technology enabled mechanistic discovery of OPC competition

Because OPC competition occurs between TN-null and wildtype OPCs, phosphoproteomic profiling to identify its underlying signaling pathways requires a platform that avoids disrupting cell-cell interactions. To satisfy this technical requirement, we adopted an “in-tissue” phosphoproteomic profiling technology that is exceptionally sensitive, with the capability to capture phosphorylation events from a few thousand cells in fixed tissues ^52^. To sift out OPC competition-related signals from the background noise of other cell types in the brain, we took advantage of the progressive nature of OPC competition in MADM-TN, which starts around P20 (only a few TN-null OPCs), peaks around P35 with ample interactions between expanding TN-null OPCs and surrounding normal OPCs, and ends around P60 as TN-null OPCs populate the entire brain (Figure 1B). By profiling WT and MADM-TN brains at each of these ages, we can filter out background signals from non-OPC brain cells by normalizing the profiling data of MADM-TN brain tissues against age-matched wildtype brains. Resulting phosphopeptides should fall into two distinct temporal patterns: those peaking at P35 but returning to baseline at P60 should be related to competition-specific signaling; those continuously increasing with age should be TN-null OPC-intrinsic signaling (Figure 5A). Procedurally, fixed MADM-TN brains along with age-matched wildtype brains were systematically cryosectioned. Sectioned brain slices were lysed, digested, and TMT-labelled for multiplexing. Phosphopeptides (phosphotyrosine followed by phosphoserine and phosphothreonine) were enriched for improved detection by LC-MS/MS (see Method). Phosphopeptides from MADM-TN brains at each age (P20, P35, and P60, respectively) were normalized with those signals from age-matched wildtype brains before further analysis (Figure 5B).

**Figure 5.**
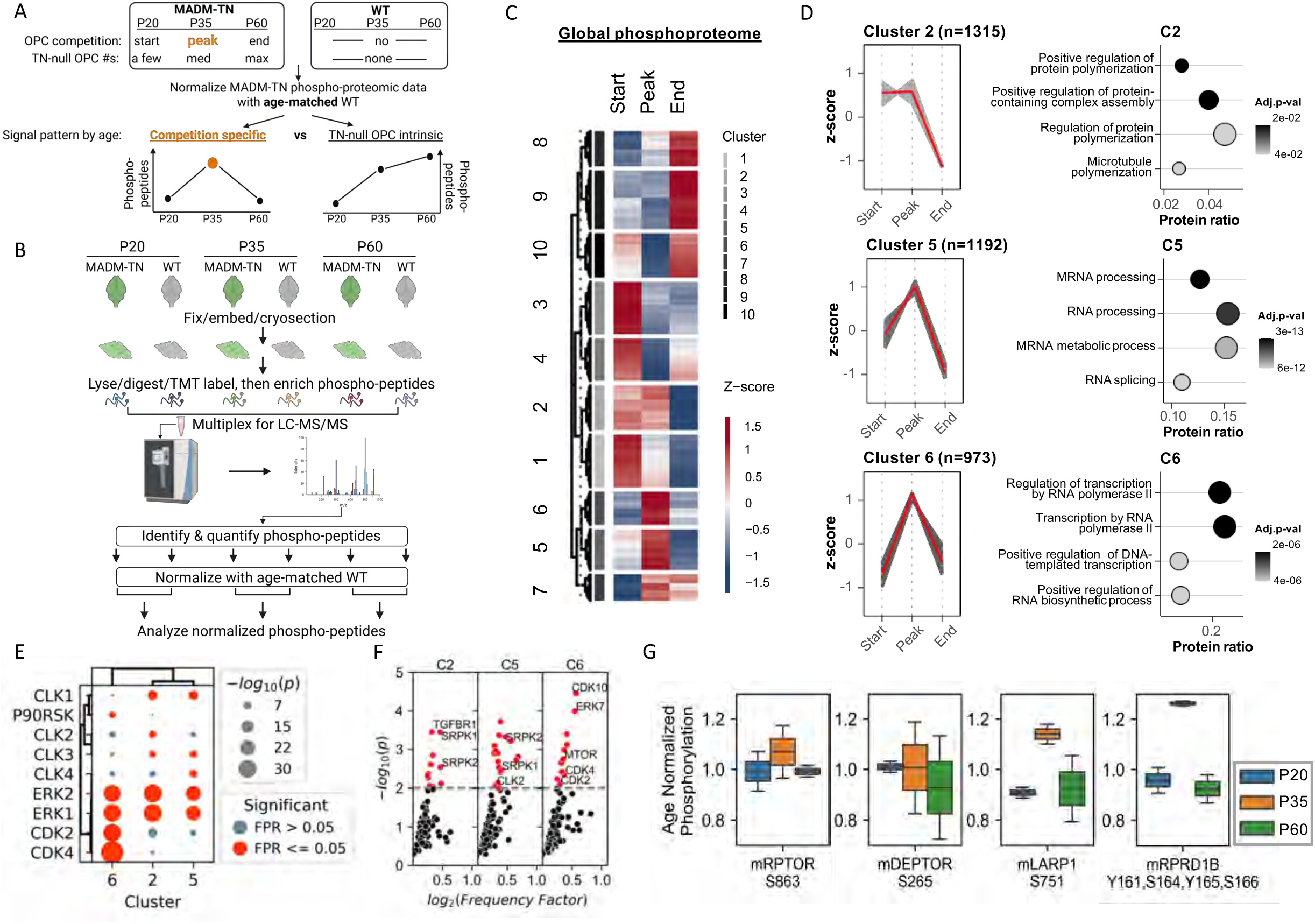
“In-tissue” phosphoproteomic profiling suggests effector mechanisms of OPC competition. **A.** Overall experimental design to distinguish competition-specific signaling from cell intrinsic signaling, based on the temporal patterns of phosphopeptides. **B.** Workflow to implement the experimental design with “in-tissue” phosphoproteomic profiling. **C.** Ten distinct clusters of co-regulated phosphopeptides identified by k-means clustering. **D.** The temporal patterns of clusters C2, C5 and C6 (left) matched the predicted patterns of competition-specific signaling, whose gene ontology analysis was performed by EnrichR (right). **E,F.** Phosphopeptides in clusters C2, C5 and C6 were further analyzed for kinase prediction, using (**E**) KSTAR and (**F**) Kinase Library methods, respectively. **G.** PhosphoSitePlus nominated mTORC1 as a candidate signaling node for cell competition.

After data processing, the normalized global phosphoproteomic (mostly serine and threonine phosphorylation) profiling at three ages revealed ten distinct clusters (C1–C10) based on k-means clustering, capturing the dynamic phosphorylation events across start, peak, and end phases (Figure 5C, Figure S10A,B). The clusters overall followed five clear temporal patterns characterized by mirroring upward and downward trends, with each cluster exhibiting distinct phosphorylation dynamics linked to coordinated modulation of functional pathways. Because OPC competition increases over time, we focused our analysis on the clusters with upward trends. Among them, C2, C5, and C6 best matched with our predicted “OPC competition” pattern (Figure 5D). Importantly, the pathways of “competition” clusters involve RNA processing and splicing, protein translation, and protein complex formation (Figure 5D), clearly distinct from the pathways involved in “intrinsic signaling” (C7, C8) (Figure S10B). These findings suggested that our profiling successfully distinguished competition-specific signaling from intrinsic signaling in TN-null OPCs. Finally, the analysis of the tyrosine phosphoproteome data revealed five distinct clusters, with C5 matching the “OPC competition” pattern (Figure S10C,D) that had an overrepresentation of cell stress-associated processes, such as p38 and HIPK1. Taken together, these data suggest that at the peak of OPC competition, TN-null OPCs generate unique phosphoproteins that confer heightened fitness to eliminate wildtype OPCs.

To identify actionable targets, we took advantage of the ability to predict which kinases are activated based on phosphoproteomic data, using the enrichment of phosphosubstrates as a proxy for kinase activity. Because phosphorylated serines and threonines are much more abundant than phosphorylated tyrosines, we focused on the former for kinase prediction, using two complementary approaches: 1) Kinase Library (Figure S11A) that has the advantage of covering all human serine/threonine kinases but only accounts for the linear motif around substrates ^63–65^; and 2) KSTAR (Figure S11B) that covers a smaller set of kinases but is informed by additional layers beyond motifs, such as protein-protein interactions ^66^. Both methods predicted activation of ERK at the peak of OPC competition, along with CDKs and kinases associated with RNA splicing regulation (Figure 5E,F). Interestingly, within clusters that peak at P35, Kinase Library predicted an enrichment of targets for mTOR for protein translation in C6 (Figure 5F), matching the prediction of protein translation by the gene ontology term enrichment (Figure 5D). Between two well-known mTOR complexes, we decided to focus on mTORC1, based on annotations from PhosphoSitePlus (Figure 5G): 1) the increased phosphorylation of Ser863 on Raptor found in C6 should facilitate the formation of mTORC1 complex; 2) the increased phosphorylation of Ser265 on Deptor found in C2 should result in its degradation and the release of its inhibition of mTORC1; 3) the increased phosphorylation of mTORC1 substrates LARP1 and RPRD1B found in C6 indicate elevated mTORC1 activity. Taken together, these data suggest that protein phosphorylation-mediated cellular signaling networks, and specifically those regulating and regulated by mTORC1, could be targeted to block OPC competition.

### mTORC1 signaling node in TN-null OPCs is specific for their competitiveness but not normal OPC biology

As our profiling nominated mTORC1 as a candidate signaling node that drives OPC competition, we first tested the mTORC1 inhibitor Temsirolimus in the *ex vivo* hippocampal slice platform. In stark contrast to the non-discriminative killing effect of MEKi, Temsirolimus specifically blocked the expansion of TN-null OPCs without detectable impact on normal OPCs in hippocampal slices (Figure S12). Encouraged by this finding, we performed a short-term *in vivo* experiment, and found that treating MADM-TN mice with Temsirolimus between P10 and P20 effectively blocked OPC competition without harming the normal OPCs (Figure S13).

While the experiments above seem promising, two important questions remain. First, because mTORC1 signaling has pleiotropic effects, including cell proliferation and survival, we wondered if the blockade of OPC competition by Temsirolimus is simply a consequence of these general effects. Second, drug treatment experiments cannot unequivocally attribute treatment effects to TN-null OPCs or other brain cells, posing a question that can only be answered by a mouse genetic model that specifically inactivates mTORC1 in TN-null OPCs. Before proceeding with the genetic approach, we first needed to make sure that mTORC1 activity is heightened in TN-null OPCs. Because mTORC1 activity in OPCs is below detection with immunostaining in brain tissues, we compared mTORC1 activity in acutely purified TN-null and wildtype (WT) OPCs and found that the phosphorylation of three mTORC1 immediate downstream targets were all significantly higher in TN-null than wildtype OPCs (Figure S14).

With this confirmation, we established a new mouse model (MADM-*Trp53,Nf1,Rptor*, referred to hereafter as MADM-TNR), in which GFP^+^ OPCs are homozygous null for *Trp53*, *Nf1*, and *Rptor* (mTORC1 subunit), herein referred as TNR-null OPCs (Figure 6A, Figure S15). A time course analysis of these mice from P10 to P60 revealed that TNR-null OPCs failed to expand their population at the gross level (Figure 6B). However, it is still possible that, rather than failing to compete, the lack of clonal expansion may be simply caused by the inability of TNR-null OPCs to survive, to proliferate, or to maintain cell fate. Our further analysis ruled out all these possibilities: first, TNR-null OPCs actually went through a transient clonal expansion by P10 to an extent that is indistinguishable to TN-null OPCs but failed to sustain as mice aged (Figure 6C,D); second, the proliferative rate of TNR-null OPCs was as high as TN-null OPCs at P20, suggesting that Raptor is not required for OPCs to divide, and declined proliferation of TNR-null OPCs at later ages is likely a consequence of their failure to compete (Figure 6E,F); third, while the clonal expansion of TNR-null OPCs stalled at P10, they maintained their OPC fate based on PDGFRα and NG2 staining (Figure 6G,H). To further ensure that mTORC1 inactivation does not cause general biological defects in OPCs, we established a MADM-*Rptor* model, in which GFP^+^ OPCs are *Rptor*-null while RFP^+^ OPCs are WT (Figure S16A). The fact that G/R ratio remained around 1 (Figure S16B, C) and no obvious defects were observed suggests that mTORC1 plays limited roles in normal OPC biology but is specifically needed for OPC competition. Finally, we thoroughly examined > 10 MADM-TNR mice long after the tumor latency age and never found any tumors, suggesting that blocking mTORC1 as a competition-specific signaling can effectively prevent gliomagenesis (Figure 6I,J). Taken together, although mTORC1 plays pleiotropic roles in other biological contexts, our data demonstrated that it is specifically required for OPC competition in the context of this study.

**Figure 6.**
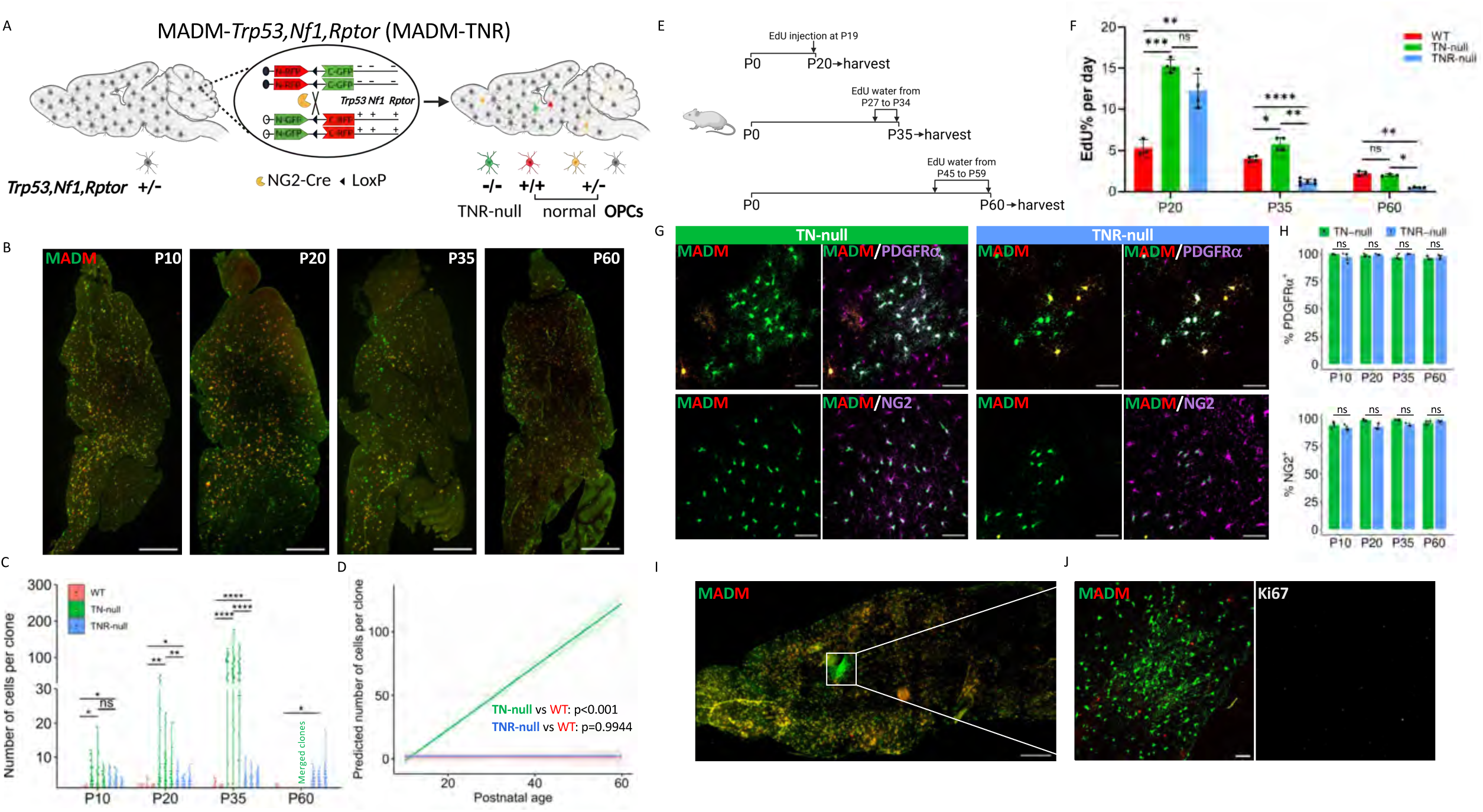
Loss of mTORC1 activity specifically in TN-null OPCs causes loss of competitiveness. **A.** Genotype-color correlation of OPCs in MADM-*Trp53,Nf1,Rptor* (MADM-TNR) brains. **B.** Macroscopic images of sagittal sections of MADM-TNR brains at various ages. Scale bar: 1,000 µm. **C.** Distribution of GFP^+^ clonal sizes in MADM-WT, MADM-TN, and MADM-TNR at various ages. 3 brains per group were used for quantification. **D.** Predicted GFP^+^ cells per clone in MADM-WT, MADM-TN, and MADM-TNR at various ages, based on a linear mixed effects model. Lines show model-predicted means; shaded areas represent 95% confidence intervals. **E.** EdU treatment scheme for measuring OPC proliferation. **F.** Proliferative rate of TNR-null OPCs at various ages, compared to TN-null and WT OPCs. **G.** Representative images showing the OPC identity of TNR-null cells (PDGFRα^+^ and NG2^+^), same as TN-null cells. Scale bar: 50 μm. **H.** Quantification of percentage of TNR-null and TN-null cells that expressed OPC markers at various ages. 3 brains per group per age were used for quantification. No significant differences were observed (p>0.05, t-test). **I.** Representative images of MADK-TNR brain at P240, the latency age of MADM-TN mice. No tumors were observed in 11 out of 11 MADM-TNR brains. Scale bar: 1000 μm. **J.** Occasional presence of GFP^+^ clusters were Ki67^-^, suggesting that these GFP^+^ TNR-null cells were not malignantly transformed. Scale bar: 50 μm.

### Relevance of OPC competition to human glioma

While our data above demonstrated the critical importance of OPC competition in pre-malignant and malignant progression, all studies were focused on *Nf1* loss-mediated OPC competition. Therefore, our final questions were: does OPC competition happen in glioma patients, and if yes, could it be driven by other gliomagenic mutations? To answer the first question, we hypothesized that, among all brain cell types, wildtype OPCs but not other cells would be specifically missing within tumor regions in patient samples. To test this hypothesis, MRI guided biopsies were obtained during gross total resection of IDH1-wildtype primary glioblastomas in 5 patients (Figure 7A) as described in Methods. Paired samples were acquired from both the solid core and infiltrating regions (Figure 7B) for single nucleus RNA-sequencing (snRNA-seq) (Figure 7C). To assess microenvironmental make-up, we calculated cell type proportions for each sample in both solid core and infiltrating regions (Figure 7D). This demonstrated an enrichment of myeloid lineage cells in the solid core as previously reported ^67^. Oligodendrocytes and, to a lesser extent, neurons could be found in both solid and infiltrating regions, with the paucity of neurons likely related to the largely white matter location of the tumors. Strikingly, there was a complete lack of OPCs in all the solid cores and in 4 of the 7 samples from the infiltrating region (we obtained two infiltrating samples from two of the five patients, and one each from the remaining three). Taken together, these findings from patient samples are consistent with the OPC competition hypothesis.

**Figure 7.**
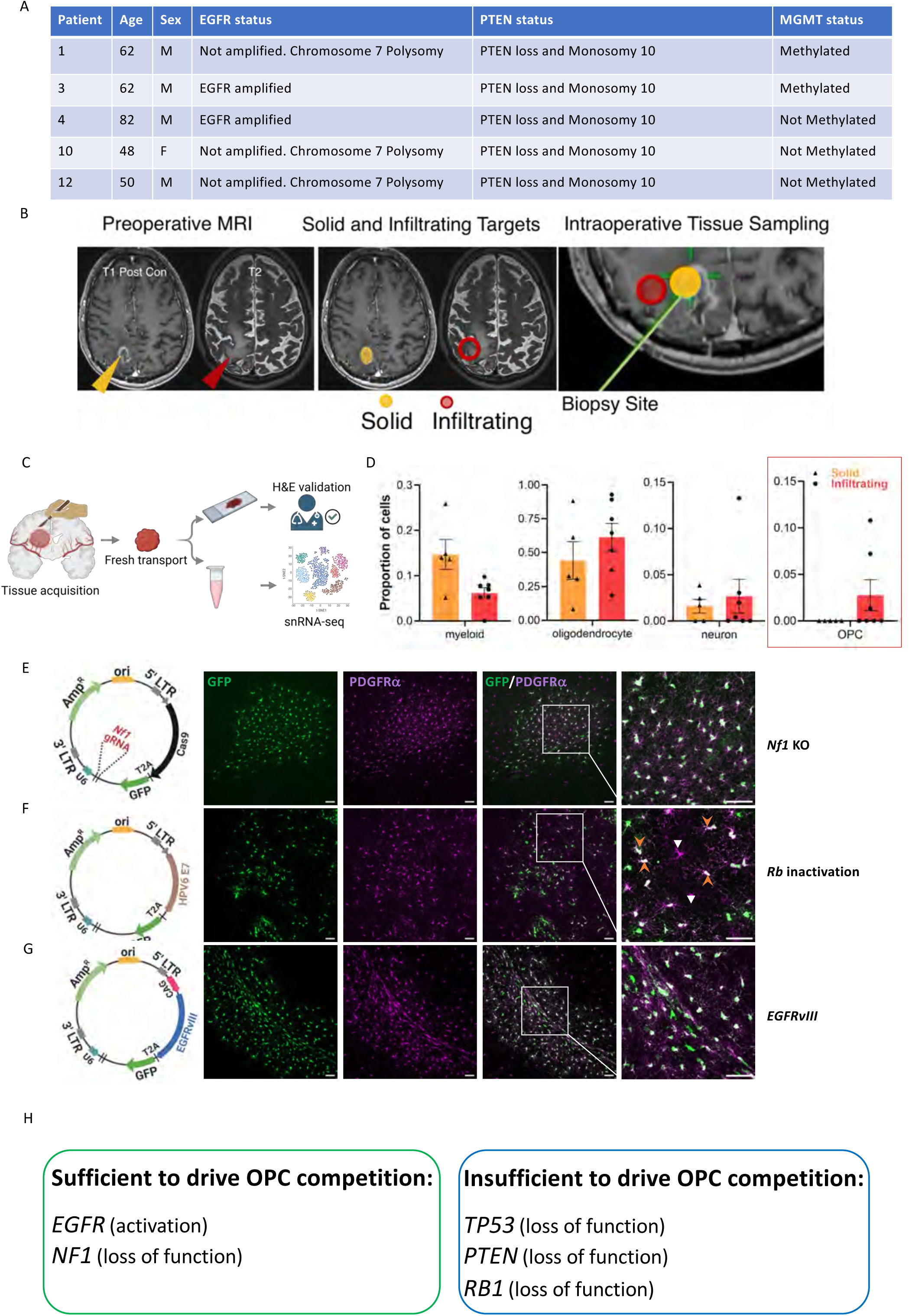
Evidence of OPC competition in adult IDH-WT glioma. **A.** Table of patient samples used in snRNA-seq, with the status of a few gene alterations listed. **B.** Sample MRIs from one patient showing T1 contrast-enhanced and T2 weighted images. On the left is the preoperative study demonstrating location of contrast enhancing zone (yellow arrow) containing the presumed solid tumor and the broader, T2-hyperintense signal (red arrow) that containing the presumed infiltrating tumor. The middle image shows the pre-operative regions selected for biopsy and the right image shows the intraoperative neuronavigation biopsy acquisition. **C.** The workflow from sample biopsy, processing, snRNA-sequencing, and data analysis. **D.** Bar plots of cell type proportions in the infiltrating and solid tumor regions. Each bar represents the mean proportion of the selected cell type in the region and the positive and negative standard deviation are represented by the bracket. Each dot represents the cell type proportion of a single sample. **E.** Representative images of expanded *Nf1-*knockout OPCs (GFP^+^PDGFRα^+^) generated by transduction of MSCV carrying *Nf1* guide RNA/Cas9. WT OPCs (GFP^-^PDGFRα^+^) were excluded from the expanded *Nf1*-null OPC clones, suggesting OPC competition. Scale bar: 50 μm. **F.** Representative images of expanded *Rb*-inactivated OPCs (GFP^+^PDGFRα^+^) upon transduction of MSCV carrying HPV-16 E7 protein that inactivates Rb function. Expanded GFP^+^ *Rb*-inactivated OPCs (orange arrow) intermixed with WT OPCs (GFP^-^PDGFRα^+^, white arrow-head), suggesting the lack of OPC competition. Scale bar: 50 μm **G.** Representative images showing the expansion of OPCs with *EGFR* mutation (*EGFRvIII*). The expanded clone of *EGFRvIII*-expressing OPCs excluded WT OPCs (GFP^-^PDGFRα^+^), suggesting OPC competition. Scale bar: 50 μm. **H.** Summary of the sufficiency and insufficiency in driving OPC competition of all examined gliomagenic mutations in this study.

Because patient samples contain many mutations, the experiment above could not answer the question as to whether gliomagenic mutations other than *NF1* can also drive OPC competition. To determine the role of individual mutations in OPC competition, we adopted a retroviral system (Figure 7E-G) that can inactivate or over-express candidate genes in OPCs in the mouse brain ^7,8,68^. We first validated the system with a *Nf1*-knockout retroviral construct, which generated a GFP^+^ OPC clone that excluded wildtype OPCs (Figure 7E), reminiscent of a key feature of OPC competition in the MADM-TN model (Figure 2B). Next, we tested G1 checkpoint and receptor tyrosine kinase (RTK) that are commonly altered in adult, IDH1-wildtype gliomas. Inactivating Rb with HPV E7^69^ resulted in proliferative GFP^+^ OPCs that intermixed with wildtype OPCs (Figure 7F), suggesting that over-proliferation due to G1 checkpoint disruption is not sufficient for competition. In stark contrast, the most prevalent RTK mutation in IDH1-wildtype gliomas (∼25%), EGFRvIII, promoted OPC competition, since transduction with *EGFRvIII*-expressing retrovirus vectors led to an expanded OPC (GFP^+^PDGFRa^+^) clone that excluded wildtype OPCs (Figure 7G). Finally, while RTKs can activate both Ras and PI3K/Akt branches, our data so far only addressed the former but not the latter. Because *PTEN*-loss that leads to aberrant PI3K activation is commonly found in IDH1-wildtype glioma patients and contributes to malignant progression ^7,8,68^, to clarify the relative competitiveness of *Pten*-mutated OPCs without ambiguity, we established a MADM-*Pten* model (Figure S17A)^70^ and performed similar analyses as we did with the MADM-TN model. To our surprise, *Pten*-mutated GFP^+^ OPCs did not outcompete normal OPCs in the brain (Figure S17B-F), suggesting that PI3K/Akt activity does not drive OPC competition and likely contributes to glioma malignancy in different ways. Taken together, this set of experiments demonstrated that OPC competition is broadly relevant in human gliomas, and can be mediated by aberrant Ras activation; however, TP53 loss, G1 checkpoint breakdown, and PI3K/AKT activation most likely play distinct roles that cooperate with OPC competition to promote malignant progression in gliomas (Figure 7H).

Finally, while the analyses above were all focused on IDH1-wildtype gliomas, we wondered if there are also indications of OPC competition in IDH1-mutant gliomas. This is important for three reasons: first, IDH1-mutant gliomas were recognized by the WHO as distinct entities from IDH1-wildtype gliomas in 2016, and comprise a large proportion of total glioma cases; second, many IDH1-mutant gliomas were reported to bear OPC signatures ^71–73^; and third, the availability of antibodies that specifically recognize the mutant form of IDH1 [MAB0733] ^74^ provides an opportunity for us to directly visualize the relative densities and distributions of glioma cells and OPCs in patient samples as a means of inferring cell competition in each tumor. Because tumor border regions are often not resected surgically in order to preserve critical neurological functions, we reviewed over 100 cases to eventually identify six samples containing distinguishable core, border, and sometimes even "normal" regions (Figure S18A-C, Table S1). Next, we attempted to distinguish glioma cells from normal OPCs with the co-staining of mutant IDH1 and olig2 (Figure S18D), since glioma cells would be expected to co-express both, while normal OPCs would similarly be expected to express only olig2. Even though there is the caveat that mutant IDH1 immunoreactivity is not always evident by immunocytochemistry, we predicted that, if competition exists in human patients, the density of olig2^+^-only cells in the tumor border would be higher than the tumor core, with a positive trend as their distance from the core increases. Because these samples had varied size of border regions, to ensure the robustness of our analysis, whenever possible, we used border regions as far as possible from the tumor core. Among six samples, other than those with complications post-treatment relapse or leptomeningeal invasion (Figure S18C, case #5 and #6), four of them showed evidence of cell competition, with more olig2^+^-only cells in the border than the tumor core (representative images and quantification in Figure S18D,E). Furthermore, in patient samples with large border regions, we noticed a trend of increased OPC density in border regions further away from the tumor core (Figure S18F, case #1 and #2). Therefore, it appears that OPC competition also occurs in some IDH1-mutant gliomas.

## DISCUSSION

In this study, we have examined gliomagenesis through the new lens of cell competition biology. We not only demonstrated the indispensable role of cell competition between mutated gliomagenic cells and non-transformed counterparts with a mouse genetic mosaic model, but also uncovered signaling mechanisms behind OPC competition with an “in-tissue” phosphoproteomic profiling platform. Functionally, we demonstrated that gliomagenesis can be effectively blocked through “power balancing” strategies, either by strengthening of “loser” OPCs by introducing the competitive mutation, or by the inhibition of competition-specific signaling in “winner” OPCs. Finally, we ascertained the relevance of OPC competition to human gliomas through both demonstrating the unique paucity of OPCs in patient samples containing high numbers of glioma cells and determining competition-conferring activities of a list of common gliomagenic mutations.

### Gliomagenic OPCs overcome contact inhibition by eliminating their normal counterparts

OPCs comprise the largest progenitor pool in the brain that maintains self-renewal in the adult brain ^37–41,44,45^. OPCs form a tiling pattern, whose mutual contacts have been found to be critical in maintaining their homeostasis. For example, the proliferation of OPCs is tightly regulated through contact inhibition ^46,47^, meaning that OPCs only enter the cell cycle when they detect a nearby void due to the death or differentiation of a neighboring OPC. Supporting this notion, during human brain development, loss of PCDH15, which typically mediates daughter OPC repulsion following a division event, led to reduced proliferation due to increased daughter cell proximity ^75^. Notably, while this study identified the molecular player behind daughter cells repelling one another after division, the proximity-triggered inhibitory mechanism is yet to be uncovered. On top of these, our discovery of OPC competition in gliomagenesis added another layer of complexity of OPC-OPC interactions, manifested as unidirectional aggression rather than mutual inhibition. This is reminiscent of recent studies on human OPCs showing that young OPCs outcompete older ones, in which age-related fitness differences provided the basis for competition ^76^. Therefore, competition between OPC populations acts as a double-edged sword, allowing regenerative cell replacement in the aging context while mediating gliomagenesis in the setting of acquired mutations favoring competitive dominance.

One of the critical questions for cell competition is its relationship with cell proliferation: which is the cause, and which is the consequence? Multiple lines of evidence in our study suggest that OPC competition enables proliferation, rather than the other way around. First, while proliferation is obviously necessary for OPC competition, we showed that it is not sufficient because *Rb*-blockade induced over-proliferation resulted in “peaceful” intermixing between mutant and wildtype OPCs. Second, the proliferative rate of TNR-null OPCs is as high as TN-null OPCs initially, whose decrease at a later stage is most likely the consequence of the inability of TNR-null OPCs to compete. Finally, while the proliferative rate of TN-null OPCs is much higher than WT OPCs during the period of competition, it becomes indistinguishable from WT OPCs at P60 as competition subsides (Figure 6F). Taken together, proliferation appears to be the consequence of competition during pre-malignant progression.

Still, there are many outstanding questions that warrant further investigation. First, is OPC competition mediated by direct contact between winner and loser cells, or by competing for limited factors or nutrients in the brain? Second, how do loser OPCs die, and how are their debris disposed of and by which cell type(s)? Third, is OPC competition a deviation of contact inhibition mechanism (uni- versus bi-directional inhibition), or an entirely novel mechanism? Finally, are mutation-mediated and age-dependent ^76^ OPC competitions mediated by identical or distinct mechanisms?

### Mechanistic insight into OPC competition

While cell intrinsic signaling has been studied broadly, cell competition signaling is understudied due to multiple technical challenges. First, winner and loser cells belong to the same cell type, making it difficult to parse out competition-specific signals from nearly identical intrinsic signaling networks. Second, competition signaling can easily get disrupted during tissue dissociation, making the workflow of cell purification detrimental for identifying competition-specific signaling. Our study was only made possible because of the adoption of the state-of-the-art “in-tissue” phosphoproteomic profiling platform.

Comparing our study with literature reports on cell competition demonstrated that cell competition signaling is highly context dependent. While *Trp53* loss has been shown to confer a winning fate during mouse embryogenesis ^77^, it doesn’t play a role in OPC competition.

Furthermore, our discovery of Ras-mTORC1 axis as the mechanism of OPC competition in our experimental models provides a new perspective of this signaling pathway. While Ras is best known for its role in driving uncontrollable proliferation in a cell intrinsic fashion, its role in aggression toward sibling cells is novel. In fact, Ras activation in epithelial cells leads to loser fate rather than winner fate, i.e. cells with over-activated Ras get extruded from epithelial sheet^78^. As importantly, while Ras-induced proliferation certainly plays an integral role in OPC competition, it appears not to be a driving force as discussed above. Based on our profiling study, it seems that the specific role of Ras signaling in the context of OPC competition is the generation of novel protein species through combined actions of regulated mRNA splicing by SRPKs/CLK2 and adaptive translation of specific protein species by mTORC1. While previous reports demonstrated the sufficiency of mTORC1 in cell competition ^79–81^, our study provided unique insights by demonstrating its necessity in OPC competition (Figure 6). Interestingly, *Pten* loss that is known to activate PI3K/Akt/mTOR signaling axis did not lead to OPC competition (Figure S17), suggesting that either mTORC1 activation is insufficient for OPC competition, or *Pten* loss failed to sufficiently activate mTORC1 in this context.

There are still remaining questions concerning the molecular mechanisms of OPC competition. First, since OPC competition is homotypic (only happens among themselves), how do OPCs recognize each other as sibling cells while ignoring other brain cell types? Second, how do winner and loser OPCs compare their fitness? Third, we still don’t know how heightened mTORC1 activity in mutant OPCs confers competitiveness: through increased expression of certain receptors to compete for rate-limited survival factors in the brain, or through generating “aggressor genes” on cell membrane to attack wildtype OPCs, or through other unknown mechanisms? Because these mechanisms likely involve membrane or secreted proteins that are presumably harder to detect than amplified intracellular signaling transducers, adopting other profiling technologies with even higher signal-to-noise ratio than the one used in this study, such as newly developed INSIGHT ^82^, would be critical.

### How could the discovery of OPC competition impact glioma treatment strategy?

The translational potential of our findings is dependent on the prevalence of OPC competition in glioma patients. This is an intrinsically difficult question not only because few have looked but also because the absence of OPC in the tumor mass could have simply been caused by false negatives due to technical problems. In this study, a carefully laid out sample collection procedure ^67^, an experimental snRNAseq scheme that included enough tumor microenvironment (TME) cells, and a meticulous analytical pipeline to clearly distinguish glioma cells from normal OPCs to avoid false calling of glioma cells as OPCs ^83^, enabled us to demonstrate the specific lack of normal OPCs but not other TME cell types in the tumor mass, implying OPC competition in human patients. To understand the genetic underpinning of OPC competition in patients, we used a medium throughput retroviral system to examine commonly altered pathways in IDH1-wildtype adult gliomas. Along with MADM-based analysis, we discovered that OPC competition is specifically mediated by *Nf1*-loss and *EGFRvIII*, but not by *Trp53* mutation, G1 checkpoint disruption, or even *Pten*-loss. Therefore, we conclude that cell competition is relatively prevalent in human gliomas, synergizing with other genetic aberrations to fuel the extraordinary malignancy.

How could the discovery of OPC competition impact clinical practices in the future? While conventional therapies focus on killing tumor cells, glioma treatment in the context of cell competition must shift the attention toward “power balancing”, through either the specific weakening of the competitiveness of glioma cells, or the strengthening of wildtype OPCs.

For the former, our study took advantage of an “in-tissue” phosphoproteomic profiling platform to nominate mTORC1 as a competition-specific candidate target, and subsequently validated its role with both pharmacological and genetic tools. Furthermore, the MADM-TNR genetic model unequivocally demonstrated that the contribution of elevated mTORC1 activity to OPC competition lies within mutant OPCs rather than other cell types in the brain microenvironment. Finally, both pharmacological experiments and MADM-*Rptor* clearly indicated that the decrease and even the loss of mTORC1 activity resulted in minimal adverse effects on wildtype OPCs.

Therefore, targeting mTORC1 can truly be considered a specific anti-competition treatment because it weakens the competitiveness of mutant OPCs without harming WT OPCs. Our finding echoes well with clinical findings. Although the initial setbacks of mTORC1 inhibitors for treating high grade gliomas were attributed to PK/PD problems ^84,85^ or rapid development of resistance ^86^, the efficacy of an mTORC1 inhibitor in LGG clinical trials clearly demonstrated the promise of mTORC1 inhibitors in glioma treatment ^87,88^. Importantly, a recent N2M2/NOA-20 Umbrella Trial showed clear clinical activity of Temsirolimus in patients whose tumors exhibited phospho-mTOR positivity ^89,90^. Therefore, while there is still debate about the efficacy of mTORC1 inhibition in glioma, our data support the idea of furthering clinical investigation, especially in combination with other agents that target the G1 checkpoint, P53 and/or PI3K pathways.

For the latter, our proof-of-concept MADM-AC and tumor grafting experiments demonstrated that strengthening normal OPCs by knocking out *Nf1*, a tumor suppressor gene, can be highly effective in preventing not only gliomagenesis but also malignant expansion. While counter-intuitive, the concept of strengthening of wildtype cells to block cancers driven by cell competition has been demonstrated by multiple studies performed in *Drosophila* models, in which apoptotic inhibitors were shown to restrain malignant expansion ^91,92^. Similarly, in a mouse model for liver cancer, the activation of YAP/TAZ oncogenes in peritumoral normal cells led to regression of tumors ^93^. Finally, mutant clones were found to outcompete emerging esophageal tumors in a mouse model, serving a tumor suppressing role ^94^. Because inactivating tumor suppressor gene *NF1* in OPCs to fight off malignant glioma cells carries risks on its own, it would be imperative to thoroughly investigate underlying mechanisms of OPC competition to enable the development of precise, competition-mitigating strategies to treat gliomas effectively.

Our study, along with the discovery of cell competition in colon cancers ^95–97^, should raise the awareness of this previously underappreciated hallmark of cancer. In models of colon cancer, it was found that either sequestering the competitive protein NOTUM from mutant cells ^95^ or the activation of Wnt signaling with lithium chloride in normal cells ^97^ could restrain the expansion of pre-malignant cells. Collectively, these studies suggest that, for cancers involving cell-competition mechanisms, effective treatment must consider both cell-intrinsic and cell-competition signaling ^98^. In particular, sibling cells acting as a unique component of TME in the context of cell competition must be taken into careful consideration for their fundamental impact on the trajectory of tumor initiation and progression and can be harnessed for the development of unconventional therapeutic strategies.

## Limitations of our studies

● While glioma cells are known to have multiple states, our studies are mostly applicable to those with the OPC-like state. Notably, literature reports on the significant presence of invasive OPC-like glioma cells at tumor margins ^32,35,36^ warrant broader investigation to determine the prevalence of OPC competition in human glioma patients.
● Our study avoided using patient glioma grafting approach to assess the human relevance of OPC competition because cross-species transplanted normal human OPCs can already outcompete rodent OPCs in the host brain ^99^, making such experiments hard to interpret - are rodent OPCs outcompeted by human glioma cells due to their gliomagenic mutations, or simply due to cross-species competition? A rodent model with humanized wildtype OPCs, once available, would be a suitable experimental platform for patient glioma grafting experiments.

## Author contributions (CRediT)

- Jiang: Conceptualization, Investigation, Formal analysis, Project Administration, Supervision, Writing – Original Draft Preparation
- Ahn: Investigation, Formal analysis, Visualization, Writing – Original Draft Preparation
- Huang: Investigation, Formal analysis, Visualization
- Gonzalez: Conceptualization, Investigation
- Kim: Investigation
- Zhang: Investigation
- Liu: Investigation
- He: Resources
- Dudley: Conceptualization, Methodology, Formal analysis, Investigation, Writing-original draft,
- Patel: Conceptualization, Resources, Methodology (surgical), Investigation, Writing-Review and Editing, Visualization
- Dzhivhuho: Methods
- Crowl: Formal analysis
- Przanowski: Formal analysis
- Quesada Camacho: Investigation, Formal analysis
- Hao: Formal analysis, Visualization
- Zeng: Methodology
- Epstein: Resources
- Hippenmeyer: Resources
- Fallahi-Sichani: Supervision
- Janes: Supervision
- Naegle: Supervision
- Hammarskjold: Supervision
- Goldman: Conceptualization, Funding acquisition, Writing -review and editing
- Kornblum: Conceptualization, Resources, Writing -review and editing, Supervision, Funding acquisition, Visualization
- Yao: Conceptualization, Funding acquisition, Project Administration, Supervision
- White: Conceptualization, Funding acquisition, Writing -review and editing, Supervision
- Zong: Conceptualization, Funding acquisition, Project Administration, Supervision, Writing – Original Draft Preparation, Writing -review and editing

## Specific figure panels or methods or reagents

- Jiang: under the supervision of Zong, designed and performed all experiments except for those specified by other co-authors
- Ahn: under the supervision of White, designed and performed the “in-tissue” phosphoproteomic profiling experiment in collaboration with Jiang and Zong, and then performed data analyses in collaboration with Crowl and Naegle for Figure 5, S10, S11
- Huang: under the supervision of Zong, performed data analysis for Figure 4M-R; performed experiment and data analysis for Figure 6C-G
- Gonzalez: under the supervision of Zong, performed experiment and data analysis for Figure 3G-L
- Kim: under the supervision of Zong, performed experiment and data analysis for Figures S3; S13
- Zhang: under the supervision of Zong, performed experiment and data analysis for Figure 1F
- Liu, He: under the supervision of Yao, collected and screened patient samples, then performed experiment and data analysis for Figure S18
- Dudley, Patel, Goldman, Kornblum: performed experiment and data analysis for Figure 7 A-D
- Dzhivhuho, Hammarskjold: provided MSCV and technical support for Figure 7 E-H
- Przanowski and Janes: data plotting for Figure 1F; modeling for Figure 2E; data analysis for Figures S3D, S9, and S12
- Quesada Camacho and Fallahi-Sichani: performed experiment and data analysis for Figure S14
- Hao: performed data analysis for Figures 2D and S5
- Zeng: developed 3D clearing method used for Figure S5
- Hippenmeyer: provided MADM19 mouse model for Figure S17
- Yao: contributed intellectually to the entire project, obtained a large collection of patient samples, and supervised experiments for Figure S18
- White: supervised the profiling experiments for Figure 5, S10, S11
- Zong: conceptualized and supervised the entire project, including the management of collaborations

## DATA AVAILABILITY

The mass spectrometry proteomics data have been deposited to the ProteomeXchange Consortium via the PRIDE [1] partner repository with the dataset identifier PXD068601 and 10.6019/PXD068601. [1] Perez-Riverol Y, Bandla C, Kundu DJ, Kamatchinathan S, Bai J, Hewapathirana S, John NS, Prakash A, Walzer M, Wang S, Vizcaíno JA. The PRIDE database at 20 years: 2025 update. Nucleic Acids Res. 2025 Jan 6;53(D1):D543-D553. doi: 10.1093/nar/gkae1011. (PubMed ID: 39494541).

## ACKNOWLEDGEMENTS

We thank Dr. Wenjie Liu for providing critical feedback on the manuscript. We also thank Dr. Pat Pramoonjago at the Biorepository and Tissue Research Facility, and Hope Davis at the vivarium for their assistance on the project. These Core Facilities are supported by UVA Cancer Center grant #P30-CA044579. We are grateful to Dr. Jonathan A. Epstein for providing the *Nf1^GRD/+^* mouse strain (https://pubmed.ncbi.nlm.nih.gov/26460546/). This work was partly supported by the National Institute of Neurological Diseases and Stroke R21 NS125479-01A1 (H.Z.), American Cancer Society Institutional Research Grant to the University of Virginia (Y.J.), the National Natural Science Foundation of China #82072787 (M.Y.), the National Cancer Institute U54 CA238114 (F.W.), U01 CA284193 (K.M.N.), and U54 CA274499 (K.A.J., M.F-S.), the National institute of General Medical Sciences R35 GM133404 (M.F-S.), the Dr. Miriam and Sheldon G. Adelson Medical Research Foundation (H.I.K., S.A.G.), the National Center for Advancing Translational Sciences KL2TR001882 (K.S.P.), Tower Cancer Career Development Grant (K.S.P.), McKnight Neurobiology of Brain Disorders Grant (K.S.P.). The content is solely the responsibility of the authors and does not necessarily represent the official views of the National Institutes of Health. Illustrations in this manuscript were created with BioRender (BioRender.com)

## Methods and Materials

### Mice

MADM-ML pair TG11ML, GT11ML JAX# 030578 ^100^, *Trp53*-KO JAX# 002101 ^101^, *Nf1*-flox

JAX#017639 ^102^, NG2-Cre JAX# 008533 ^103^, *Trp53*-flox JAX# 008462 ^104^, *Rptor*-flox

JAX#013188 ^105,106^, *Pten*-flox JAX#006440 ^107^, NG2-CreER JAX#008538 ^108^, *hGFAP*-Cre

JAX# 004600 ^109^, *Rosa26-LSL-tdTomato* JAX# 007908 ^110^ were requested to be used in this study. Nude (Hsd:Athymic Nude-Foxn1) mice were purchased from Envigo.

MADM pair on chromosome 19 ^70^ are accessible at the European Mouse Mutant Archive (EMMA) with the following RRID and information: MADM-19-GT - https://www.infrafrontier.eu/emma/strain-details/?q=14720 -RRID:IMSR_EM:14720; MADM-19-TG - https://www.infrafrontier.eu/emma/strain-details/?q=14721 -RRID:IMSR_EM:14721.

*Nf1^GRD/+^* mouse line ^111^ was provided by the Jonathan A. Epstein lab at the University of Pennsylvania, who worked with GenOway to use an elegantly designed ES targeting strategy to replace exon 28 of *Nf1* with a R1276P point mutation, selected *in vitro* with a Neomycin cassette embedded in intron 27. After verifying homologous recombination in ES cell clones, blastocyst injection, germline transmission, and Cre-mediated excision of Neomycin cassette, the modified *Nf1* locus in resulting mice is identical to WT allele except for a loxP site in intron 27 and a R1276P point mutation in exon 28. For genotyping, primer set of Forward-5’ GGG GCT TCT GGG GTA TTT AT 3’ Reverse-5’ CAA GAC ACT GGA GTA AGG GAA AA 3’ was used. A 61bp band indicates the WT allele while the mutant allele can be identified with a 134bp band.

For all *in vivo* experiments, the ages of mice were indicated in figures and/or figure legends. Mice were separated into experimental groups based on genotype and age. Similar number of female and male mice were used in all experimental groups to control for sex-related effects. All animal procedures were performed according to the protocols approved by the Institutional Animal Care and Use Committee (IACUC) of the University of Virginia. Statistical tests were used to predetermine sample size of experimental mice.

### Brain tissue preparation and histology

We performed the brain tissue harvest procedures as described previously ^9^. In brief, mice were anaesthetized (60mg/kg ketamine; 12 mg/kg xylazine) and perfused through the left cardiac ventricle with PBS, followed by 4% paraformaldehyde (PFA). Brains were then post-fixed overnight at 4°C, cryoprotected in 30% sucrose (two days at 4°C) and embedded into optimal cutting temperature (O.C.T.) prior to cryo-sectioning on a cryostat. 16-25 μm sections were captured onto plus slides and dried at room temperature for 1 hour before proceeding to immunohistochemistry.

### 5-ethynyl-2’-deoxyuridine (EdU) labeling and detection

EdU (Invitrogen) was administrated by intraperitoneal injection at 10mg per kilogram body weight or by drinking water at 0.2mg/ml for long term labeling (7-14 days). Mice were sacrificed right after the treatment and brain tissue was collected. EdU positivity was visualized by CLICK-IT reaction using AlexaFluor 647 conjugated Azide. Dried sections were re-hydrated with PBS, permeabilized in PBS/0.3% (v/v) Triton X-100 for 1 hour and then transferred to the EdU developing cocktail, incubated in the dark for overnight. Then the slides were washed free from developing reagents and proceed to standard immunohistochemistry.

### Immunofluorescence staining and imaging

We followed the standard procedures as described previously ^9,112^. Briefly, dried brain sections were rehydrated with PBS, blocked in 5% normal donkey serum (NDS) in PBST (PBS + 0.3% Triton X) for 30 minutes at room temperature to prevent non-specific binding. Then the sections were incubated with primary antibodies (diluted in 5% NDS/PBST) overnight at 4°C. After several washes with PBST, appropriate fluorophore conjugated secondary antibodies (donkey as host, Jackson ImmunoResearch) were applied at 1:250 for overnight at 4°C or 4 hours at room temperature. Following several washes with PBST, the sections were stained with DAPI (1ng/ml in PBS) to visualize the nuclei. Finally, slides containing the brain sections were mounted with anti-fade mounting gel mounting medium (Electron Microscopy Sciences).

In the cases that EdU visualization occupied the far-red (Alexa 647) channel, PDGFRα staining would occupy the UV-channel which was usually used for visualizing nuclei by DAPI. An appropriate biotin-conjugated secondary antibody would be used to recognize the primary antibody and then Alexa Fluor 405 conjugated streptavidin to develop the fluorescent signal.

Primary antibodies used were listed below: GFP (Chicken, Aves Labs), DsRed (Rabbit, Clontech), mouse specific PDGFRα (Goat, R&D), Ki67 (Rabbit, Abcam), NeuN (Rabbit, Abcam), Olig2 (Rabbit, Millipore), Sox9 (Rabbit, Abcam), NG2 (Rabbit, Millipore).

Images were acquired on Olympus MVX10, Nikon epi-fluorescence scope or Zeiss confocal system (LSM710 or 900) and processed using Fiji software.

### Quantification of G/R ratio and total OPC number

Fixed mouse half brains were cryosectioned sagittally starting from midline. 4-5 sections (30 μm in thickness) with at least 200 μm between the sections, which cover the medial to lateral axis, were collected on a plus side and stained with GFP, RFP and PDGFRα. Images were taken from olfactory bulb, cortex, thalamus, ventral striatum and hypothalamus regions ^112^ from each section. PDGFRα^+^ was scored for total OPCs while PDGFRα^+^GFP^+^RFP^-^ was scored as *Trp53,Nf1*-null OPCs and PDGFRα^+^GFP^-^RFP^+^ was scored as red OPCs. To take the medial/lateral difference into account, we averaged the numbers of all collected sections across medial/lateral axis. G/R ratio was calculated by dividing the number of *Trp53,Nf1*-null OPCs by that of red OPCs. Total OPC numbers were calculated as the average of PDGFRα^+^ across all sections.

### Quantification of absolute normal OPC number of the whole mouse brain

Fixed mouse half brains were sectioned mediolaterally, and one every 30 sections were collected (30 μm thick). Images were taken in all brain regions including olfactory bulb; frontal, middle and rear cortex; thalamus; caudate putamen; ventral striatum; hypothalamus; midbrain; brain stem and cerebellum regions. Total OPCs (PDGFRα^+^) and *Trp53,Nf1*-null OPCs (PDGFRα^+^GFP^+^RFP^-^) were counted for each section. To calculate the OPC numbers of the whole brain, we first assumed that the section we collected represented the 29 sections that were skipped. Secondarily, we assumed that the two sagittaly cut half brains were the same. To calculate the total OPCs of the whole brain, we summed up the PDGFRα^+^ scored per section (usually 6 sections could cover the half brain), multiply by 30 to get the number of half brain, then multiple by 2 for the whole brain. Similar calculation was done for *Trp53,Nf1*-null (PDGFRα^+^GFP^+^RFP^-^) OPCs. The normal OPC number was deduced by subtracting *Trp53,Nf1*-null OPC number from the total OPC number in the entire brain.

### Mutant clone imaging and quantification at 3-D

Fixed mouse half brains were cleared with CUBIC reagent as described ^113^ and applied to Light Sheet Microscopy. Images were processed with Imaris software.

We used Density-based Spatial Clustering of Applications with Noise (DBSCAN) algorithm for the counting of GFP^+^ clones and the number of cells of each clone. Briefly, we first used Gradient Boosted Tree (GBT) pixel classifier to remove the background in the light-sheet images of the cleared brains. Then, individual GFP^+^ cells were segmented based on intensity thresholding. The segmented cells are termed as centroids. Lastly, we performed unsupervised clustering on the centroids using the DBSCAN algorithm from the Scikit-Learn Python library. Python code and example centroids data are available on GitHub (https://github.com/szh141/UVA-AMF/blob/main/clone_count.py). Note: there are two settings that define the clonal boundaries: 1) eps: this parameter defines the distance between GFP^+^ cells to be called into a clone. Due to neonatal (P15) brains are relatively smaller than adult (P21, P28) brains, we set eps 65 for P15 while 75 for P21 and P28 brains; 2) min_points (minPts): this parameter defines the minimum number of cells per clone. As clones grow bigger, minPts should be adjusted so that large clones will be segmented correctly. For P15, we set minPts as 5; P21 as 10 and P28 as 15. The parameters were verified by visually comparing the centroids with light-sheet images (after background subtraction) before proceeding to automatic cell/clone counting with the program.

### Mathematical modeling of clonal expansion of mutant OPCs

To evaluate the characteristics of clonal growth, we developed stochastic models of clone expansion from P15 to P21 and from P21 to P28, using observed clone sizes at the earlier timepoint as input. Each model was parameterized by free components:

- a: a constant baseline growth factor,
- b: the strength of size-dependent growth, where larger clones grow more (or less) than smaller ones, modeled via a log₂ dependency,
- λ: the mean of a Poisson-distributed noise term introducing stochastic variability in clone growth,

For a given parameter set, we quantified model fit using the Kolmogorov–Smirnov statistic between the simulated and observed clone size distributions at the later timepoint. This process was applied independently to both transitions (P15→P21 and P21→P28). To assess whether inclusion of all free parameters was justified, we compared the full model to restricted versions (with fixed b = 0), while accounting for model complexity using the Bayesian Information Criterion (BIC) [PMID 15117008]. The BIC is defined as −2 log *L* + *k* log n, where log *L* is the log-likelihood of the observed data under the model, *k* is the number of estimated parameters, and n is the number of observations. Log-likelihoods were approximated by evaluating the kernel density of simulated distributions at the observed data points. To compare models, we also computed BIC weights, which approximate the posterior probability of each model under equal priors. This allowed us to objectively assess whether incorporating size-dependent growth improved model performance.

### Tumor cell grafting

For mouse tumor cell grafting, tumor bearing (symptomatic) MADM-*Trp53,Nf1* mice were sacrificed. Tumor tissues (GFP^+^) were carefully dissected out under fluorescence microscopy (Olympus), minced into ∼1mm^3^ pieces and digested in well-equilibrated (in 5% CO_2_/95% O_2_) 1X EBSS solution with 0.5mM EDTA, 10mM HEPES, 26mM NaHCO3, 0.16mg/ml L-Cysteine, DNase and Papain (20Unit/ml) at 37°C for 45 minutes to 1 hour. Digested tissues were then washed free of Papain and briefly triturated with serological pipet in well-equilibrated 1xEBSS buffer containing ovomucoid, a trypsin inhibitor. Dissociated cell suspension was carefully layered on top of ovomucoid containing 1xEBSS solution well equilibrated in 5% CO_2_/95% O_2_ and centrifuged at 220 g for 15 minutes. Cell pellets were re-suspended in Neurobasal medium supplemented with L-glutamine, Penicillin/ Streptomycin solution, B27 supplement, PDGF-AA (10ng/ml) and NT-3 (1ng/ml). Live OPC cell number was determined by Trypan blue counting with a hemocytometer. 2-3 μL of cell suspension with density of 100,000/μL was injected into right striata of 4-6 weeks old Nude (Envigo) brains as previously described (PMID: 17296553).

For tumor grafting into hosts with different genotypes of OPCs, neonates (P4-6) with *Nf1*-null OPCs and littermates with WT OPCs were used. Tumor cell grafting was performed according to the procedure as described ^114^. Specifically, the pups were first cryo-anesthetized and 1 x 10^5^ donor cells were injected in equal amounts into the forebrain at 2 locations: 1.0 mm posterior bregma and 1.0 mm bilaterally from the midline and 1.2mm to 1.5mm depth. All cells were injected through 10 μL Hamilton Syringes inserted directly through the skull into the target sites.

Grafted mice were harvested at 5 weeks post-injection by cardiac perfusion with 4% PFA. Fixed brain tissues were dissected out and coronally cut along the needle track before proceeding to cryopreservation and sectioning.

### Proteomics and phosphoproteomics by liquid chromatography-tandem mass spectrometry (LC-MS/MS)

Whole proteomic and phosphoproteomic analyses were performed using a modified version of previously described protocols ^115,116^. Sectioned tissues were lysed in 500 μL 5% sodium dodecyl sulfate in 50 mM Tris pH 8.5 / HPLC water. Lysates were sonicated and cleared via centrifugation and prepared for mass spectrometry using a modified S-trap protocol (Protifi) version. Proteins in lysates were reduced with 10 mM Dithiothreitol for 30 min at 56°C and alkylated with 55 mM Iodoacetamide for 30 min at room temperature in the dark. The lysates were acidified to a final concentration of 2.5% v/v phosphoric acid. A 6× volume of S-trap binding buffer (90% methanol, 100 mM TEAB, pH 7.55) was mixed into each sample to precipitate proteins. The solution was sequentially loaded onto S-trap micro columns (Protifi) and spun at 4000 × g for 30 seconds until all the solution had passed through the column. The columns were washed three times with 300 μL 50% chloroform/50% methanol solution, followed by the three washes of 300 μL S-trap binding buffer. Proteins were digested on-column with 5 μg of trypsin (Promega) in 50 mM HEPES pH 8.5 overnight at 37°C in a humidified incubator. Digested peptides were eluted in three steps at 4000 × g for 1 min: 40 μL of 50 mM HEPES, 40 μL of 0.2%formic acid, and 40 μL of 50% acetonitrile/HPLC water.

Peptides for TMT-labeled analyses were lyophilized and resuspended in 50 mmol/L HEPES (pH 8.5). TMTPro 18-Plex (0.45 mg; Thermo Fisher Scientific. Lot# XI346567 and XJ346678) was resuspended in 15.3 μL of anhydrous acetonitrile, and 15 μL was subsequently added to each sample, followed by a 2-hour incubation (Supplement Table 2). Reactions were quenched with the addition of hydroxylamine to a final concentration of 0.3%, pooled, dried by vacuum centrifugation and lyophilization, and stored at –80°C before analysis. For the tyrosine phosphoproteomic analysis, the enrichment of phosphotyrosine (pTyr) peptides was performed by incubating digested peptides with 60 μl of protein G agarose beads (Sigma) pre-conjugated to 24 μg of 4G10 (Bioxcell) and 6 μL of PT66 antibodies overnight at 4 °C in immunoprecipitation (IP) buffer (100 mM Tris-HCl, 1% NP-40, pH 7.4). Subsequently, immunoprecipitated peptides were eluted twice with 0.2% trifluoroacetic acid (TFA) for 10 min each and enriched for phosphopeptides using ferric nitrilotriacetate (Fe-NTA) columns (Thermo). Eluates from the Fe-NTA columns were dried using a vacuum centrifuge, reconstituted in 3% acetonitrile/0.1% formic acid, and directly bomb-loaded onto a hand-packed 10 cm analytical column containing 3 μm C18 beads. Agilent 1100 Series HPLC connected to Orbitrap Exploris 480 Mass Spectrometer was operated at 0.2 ml/min flow rates with a precolumn split to attain nanoliter flow rates through the analytical column and nano-electrospray ionization tip. Peptides were eluted with the increasing concentrations of buffer B using the following 140-minute gradient settings (Buffer A: 0.1% acetic acid; Buffer B: 70% acetonitrile/0.1% acetic acid): 0 min: 0% B; 10 min: 11% B; 110 min: 32% B; 125 min: 60% B; 130 min: 100% B; 128 min: 100% B; 130 min: 0% B. The MS parameters were: ESI spray voltage, 2.5 kV; no sheath or auxiliary gas flow; heated capillary temperature, 275 °C, data-dependent acquisition mode with the scan range of 380–1800 m/z. MS1 scans were acquired at 120,000 resolution, maximum injection time of 50 ms, normalized AGC target of 300%, and only included the precursor charge states of ≥ 2 and ≤ 6. For every full scan, MS/MS spectra were collected during a 3-second cycle time. For MS/MS, Ions were isolated (0.4 m/z isolation window) for the maximum IT of 250 ms, normalized AGC target of 1000%, HCD collision energy of 33% at a resolution of 1200,000, and the dynamic exclusion time of 35 seconds.

For global phosphoproteome (pSer/pThr) and protein expression analyses, half of the supernatants from the pTyr-IP were subjected to high-pH reverse-phase fractionation on a Kromasil® C18 HPLC Column (5 μm particle size, pore size 100 Å, L × I.D. 250 mm × 4.6 mm) using buffer A (10 mM triethylammonium bicarbonate (TEAB), pH 8) and buffer B (10 mM TEAB, pH 8, 99% acetonitrile) over an 85-minute gradient (0 min: 1% B, 1 min: 1% B, 5 min: 5% B, 65 min: 40% B, 75 min: 70% B, 84 min: 70% B, 85 min: 1%) into 10 fractions (concatenated) using a Gilson FC 204 Fraction Collector. Fractionation was performed at a flow rate of 1ml/min and 1 min per fraction between 10-85min portion of the gradient. For each fraction, 1/10 was allocated for protein expression analysis, and 9/10 was allocated for global phosphoproteomic (pSer/Thr) analysis. Fractions for pSer/pThr analysis were resuspended in 50 μl of 0.2% TFA and subjected to Fe-NTA column-based phosphopeptides enrichment, after which they were eluted and dried. Both groups of fractions were resuspended in 3% acetonitrile/0.1% formic acid and injected into the mass spectrometry using an Orbitrap Exploris 480 Mass Spectrometer (Thermo Fisher Scientific), coupled with an UltiMate 3000 RSLC Nano LC system (Dionex), Nanospray Flex ion source (Thermo Fisher Scientific) and column oven heater (Sonation; operated at 45°C). Peptides were eluted using a 120-minute gradient (Buffer A: 0.1% formic acid, Buffer B: 80% acetonitrile/0.1% formic acid) with the following minute: B% profile: 0:3, 30:3, 32:6, 70:19, 87:29, 96:41, 99:97, 106:97, 106.1:3. The flow rate of 0.35 μl/min was used between 0-32 min and 0.1 μl/min between 30-120 min. The MS parameters were spray voltage, 2.5 kV; no sheath or auxiliary gas flow; heated capillary temperature, 275 °C; data-dependent acquisition mode with a scan range of 380–2,000 m/z. MS1 scans were acquired at 60,000 resolution, maximum injection time of 25 ms, 300% normalized AGC target, and only included the precursor charge states of ≥ 2 and ≤ 6. For every full scan, MS/MS spectra were collected during a 3-second cycle time. For MS/MS, Ions were isolated (0.4 m/z isolation window) for the maximum IT of 150 ms, normalized AGC target of 100%, HCD collision energy of 33% at a resolution of 60,000, and the dynamic exclusion time of 30 seconds.

### Proteomic and phosphoproteomic data processing

Mass spectra were analyzed using Proteome Discoverer (v3.0, Thermo Fisher Scientific) and searched using Mascot (v2.8) against the mouse Swiss-Prot database (v2023). Proteomic and phosphoproteomic data were searched with two or fewer missed cleavages, precursor and fragment ion matched with 10 ppm and 20 mmu mass tolerances, and fixed modifications of carbamidomethyl (Cys) and TMTpro (lysine and N-terminus) and dynamic modifications of oxidation (Methionine). For phosphoproteomic data, dynamic modifications for phosphorylation (Serine, Threonine, Tyrosine) were added. Peptide spectrum matches (PSMs) were filtered by an ion score ≥20 and aggregated based on unique identification. Percolator [PMID 17952086] was used in the search node for peptide identification. The ptmRS node in Proteome Discoverer was used to confidently assign the phosphorylated and oxidized sites (filtered for peptides with localization probability ≥ 90% after search). All PSMs were processed in R (v4.3.1) and R studio (v2023.12.1).

### Proteomic and phosphoproteomic data analysis

To identify the most relevant serine/threonine kinases within each cluster of the phosphoproteomic dataset, we used two main tools: Kinase Library ^63,64^ (https://kinase-library.mit.edu/ea) and KSTAR ^66^. For Kinase Library, the mouse phosphopeptides associated with a cluster were tested for the enrichment of a kinase’s motifs using the binary enrichment functions of the kinase-library python package (v1.3). The foreground consisted of all serine/threonine phosphorylation motifs identified in the given cluster and the background consisted of all serine/threonine phosphorylation motifs identified in the entire dataset. For KSTAR, as it relies on human kinase-substrate predictions from NetworKIN, the identified mouse peptides/sites were first converted to their homologous human peptides/sites using PTMoreR ^117^. The converted data was then processed and run through the KSTAR algorithm for each cluster, using every site associated with the cluster as evidence. For the enrichment analysis of Gene Ontology terms for Biological Processes (v2023) using the proteomics data, we used EnrichR ^118^.

The mass spectrometry proteomics data have been deposited to the ProteomeXchange Consortium via the PRIDE ^119^ partner repository with the dataset identifier PXD068601 and 10.6019/PXD068601.

### Brain slice culture, imaging and drug treatment

Procedures of preparing and culturing brain slices *ex vivo* were performed as described ^120^. For consistent survival, we use neonatal pups from P7 to P10 and isolate hippocampus for slice preparation. Briefly, tissue culture inserts were pre-soaked for at least two hours with slice culture media containing 50% MEM, 25% HBSS, 25% horse serum, 1mM glutamine, 25mM HEPES, 1mg/L insulin and 0.4mM ascorbic acid at pH of 7.2. Dissection buffer (124mM NaCl, 3mM KCl, 1.25mM KH_2_PO_4_, 4mM MgSO_4_, 2mM CaCl_2_, 26mM NaHCO_3_, 10mM D-(+)- Glucose, 2.0mM Ascorbic acid and 0.075mM Adenosine) was well-equilibrated in 5% CO_2_/95% O_2_ at least 15 minutes prior to the start of dissection. All tools and manual tissue chopper were sterilized with 70% ethanol. The pup was decapitated and the whole brain was isolated and place into pre-chilled and pre-equilibrated dissection buffer. Then a sagittal cut of the brain was made, and hippocampi were isolated from both hemispheres under dissection scope. The isolated hippocampi were further cut into slices of 300 μm thickness with tissue chopper. The cut slices were then transferred to a 35mm dish filled with ice-chilled oxygenated dissection buffer and separated into single slice with spatula under scope. Individual slice was then transferred with a wide-open tip and loaded onto the membrane insert pre-soaked with culture media. Each insert can accommodate 4 slices. The membrane inserts were placed on top of a thin layer of culture media in a multi-well plate. Change the culture media one hour after slice preparation and then every other day afterwards.

Allow the slices to adapt to *in vitro* culture for 5-7 days before imaging or starting drug treatment. Fluorescence images of each slice were acquired with Olympus MVX10 Macro View. Trametinib or Temsirolimus can be added directly to the culture media for treatment and replenished every other day until the end of experiment.

### Single cell measurement of mTORC1 signaling activities in cultured OPCs

*Trp53,Nf1*-null and WT OPCs were purified from neonatal pups by immunopanning as described^121^. *Trp53,Nf1*-null OPCs were purified from pups of the genotype of *p53flox,Nf1flox*/*p53flox,Nf1flox*; *hGFAP-Cre*; *LSL-fluorescent reporter*. WT OPCs were purified from pups with the genotype of *hGFAP-Cre*; *LSL-fluorescent reporter*. Acutely purified *Trp53,Nf1*-null and WT OPCs were each seeded on a PDL-coated well of a thin-glass bottom 96-well plate for 24hr before harvesting for immunofluorescent staining of mTORC1 substrates: p-S6 (Ser240/244), p-S6 (Ser235/236), and p-4EPB1 (Thr37/41), respectively. Stained cells were imaged with an Operetta CLS High-Content Imaging System. To avoid the impact of cell density on mTORC1 activity, we ensured the equal cell density at the time of harvest by confirmation with visual and Hoechst staining. Although immunopanning can yield OPCs with more than 90% purity, there are still some contaminating cells such as microglia. To ensure the analysis of mTORC1 activity only in OPCs, we first identified reporter-positive cells by thresholding the single cell mean intensity distribution: a cutoff was chosen at the trough between the two density peaks to robustly separate expressing from non-expressing cells. Using this threshold, approximately 98 % of *Trp53,Nf1*-null and 89 % of WT OPCs were identified for further analysis. Approximately 10,000 cells pooled unbiasedly from replicate wells were analyzed. The imaging data was background-subtracted and quantified using ImageJ and Cell Profiler, and the log10-trasformed single-cell distribution of protein intensities was reported in arbitrary units (a.u.).

### MSCV viral vector construction, viral packaging and injection

Murine Stem Cell Virus (MSCV) is a type of retroviral expression system engineered for stable, high-level gene delivery in dividing cells. Here we used this system to either knockout or express a gene of interest in OPCs, a population of proliferating brain cells during the neonatal age.

To knockout *Nf1* in OPCs of mouse brains, a guide sequence (5′-CAGAACAGCATCGGTGCCGT-3′) was inserted under the control of the U6 promoter in an MSCV viral vector (Addgene#124889) that also expressed Cas9 fused with EGFP via a T2A peptide. Thus, EGFP served as a reliable report of cells in which *Nf1* knockout was achieved.

To express *EGFRvIII*, a prevalent *EGFR* mutation in human glioma, the fragment was amplified from pCDNA3.1-EGFRvIII plasmid (a gift from Dr. Adrian Shimpi and Dr. Kristen Naegle) and used to replace *Cas9* in MSCV vector. The inserted EGFRvIII fragment was cloned in frame with T2A-EGFP, allowing EGFP to report EGFRvIII-expressing cells. Additionally, a CAG promoter was inserted upstream of the EGFRvIII start codon to enhance expression.

To inactivate Rb function, the HPV-16 E7 fragment was amplified from plasmid <u>p1324 HPV-16</u> <u>E7</u> (Addgene#8643) and cloned to replace *Cas9* in MSCV vector, maintaining the in-frame fusion with T2A-EGFP for accurate reporting. No exogenous promoter was added, and HPV-16 E7 expression was driven by the MSCV viral promoter.

MSCV virus particles were packaged by co-transfecting each MSCV construct with the packaging plasmid pHit ^122^ and the envelope plasmid VSV-G (Addgene #12259) into 293T/17 cells. 48hr post-transfection, viral supernatants were collected, cleared of cell debris and concentrated by ultracentrifugation (25,000rpm for 1hr) to achieve titer of about 10^7^ pfu/ml. For in vivo delivery, 1-2 μL of the viral particle suspension was injected into the brains of postnatal day 2-4 (P2-P4) wild-type mice as described ^123^. Multi-focal injections were performed to maximize transduction efficiency. Virus-injected mice were harvested at 45 days post-infection (dpi) for histological analyses.

### IDH WT patient glioma sample analysis

Biopsies: pre-operative magnetic resonance imaging (MRI) was acquired within 2 weeks of surgery and pre-processed. Thin cut (5mm) T1 with contrast sequences are used to identify contrast enhancing regions of tumor and T2 and FLAIR sequences without contrast are used to identify non-enhancing) tumor. Using segmentation software AFNI, CE and NE regions were segmented and identified using multidisciplinary (neurosurgery and radiology) consensus.

Biopsy targets were planned and uploaded to neuronavigation software (*Brainlab Curve, Munich, Germany*). During standard of care surgery, these biopsy targets were acquired prior to tumor resection to avoid brain shifts. Intraoperative re-registration was used on fixed structures (dural folds, skull base) to minimize registry inaccuracy during resection. Biopsy samples were examined by H&E staining to confirm that samples from contrast enhancing regions contained tumor cells with the characteristics of solid tumor regions having classic features of glioblastoma (high cell density, microvascular proliferation, pseudopalisading necrosis) and that nonenhancing regions contained cells with infiltrating region characteristics having tumor cells within more normal looking neuropil. If neurosurgical, pathologic review had a consensus then solid and infiltrating region samples were considered confirmed for further experiments.

SnRNA-seq: libraries were generated from nuclei using the v3 10X 3’ GEM kits. Illumina sequencing was performed on the Nova-seq machine. CellRanger was used to align and process sequencing data. Ambient RNA was removed using the CellBender program ^124^. Seurat ^125^ was utilized to normalize and cluster cells, DoubletFinder ^126^ was used to remove doublets, and Harmony was utilized to merge all samples. Tumor cells were identified using our CNV analysis pipeline with the InferCNV program, for inferred amplification of chromosome 7 and deletion of chromosome 10 ^67^. Non-transformed cell types were identified using literature search of cluster marker genes, canonical cell type markers and Azimuth cell-typing software with the human motor cortex reference ^127^. Canonical cell type markers ultimately used to identify OPCs in tumor negative cells were PDGFRA, CSPG4 and SOX10 as these met all criteria.

### IDH mutant patient sample staining and imaging processing

Tumor tissues collected from glioma patients were immediately fixed in 10% formalin and paraffin embedded with SAKURA Tissue—TEK VIP® 6 AI Vacuum Infiltration Processor and LEICA EG 1150H. Tissue sections of 3-5 μm were collected, incubated at 70℃ for 2h to ensure their attachment to the slide. The slide was then immersed in fresh Xylene for 30 and 5 minutes, followed by 100% ethanol for 5 mins twice, 95% ethanol for 5min twice, 80% ethanol for 5min, 75% ethanol for 5min, brief rinsed in and water and transferred to TBST buffer. Antigen retrieval was performed prior proceeding to immunohistochemistry. Images were obtained from Olympus BX53M fluorescence microscopy and processed with Fiji software. IDH1 antibody was from MXB Biotechnologies; olig2 antibody was from Millipore.

### Statistical analysis

Statistical significance was determined using GraphPad Prism 7 or later version using paired or unpaired Student’s two-tailed t-test, one-way ANOVA or two-way ANOVA, according to test requirements. No inclusion/exclusion criteria were pre-established. A P value <0.05 (indicated by one asterisk), <0.01 (indicated by two asterisks), <0.001 (indicated by three asterisks), <0.0001 (indicated by four asterisks) was considered significant. A P value >0.05 (indicated by ns) was considered not significant.

### Study Approval

The snRNAseq study of patient samples was approved by the UCLA IRB #10-000655. The immunohistological analysis of patient samples was approved by Sun Yat-Sen University IRB #B2021-232-01. All patients provided informed consent for all medical and surgical procedures and involvement in research studies.

---

**Figure S1.**
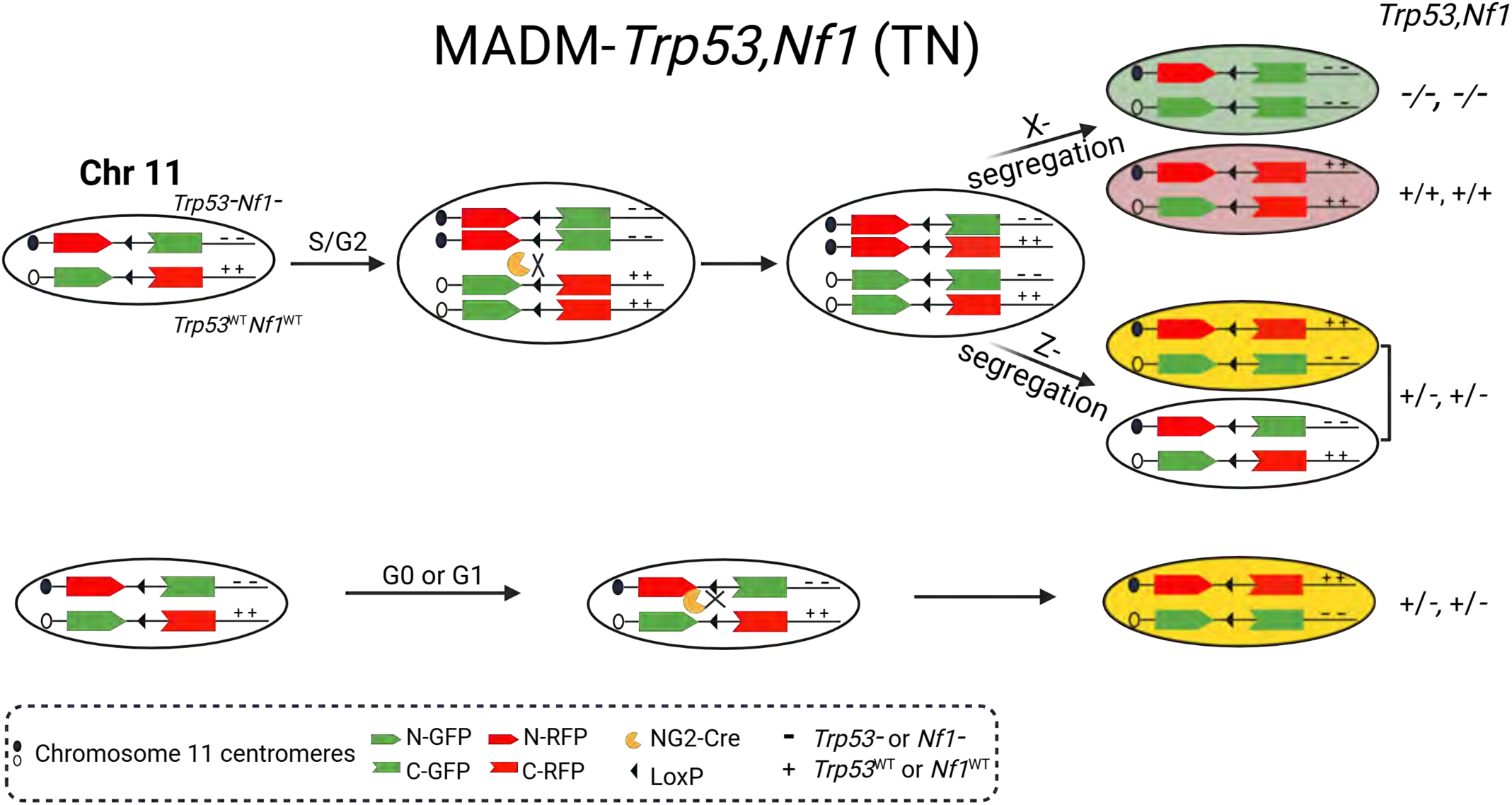
Detailed schematic of MADM-*Trp53,Nf1* (MADM-TN) model. From an otherwise heterozygous, non-labeled mouse, NG2-Cre mediated mitotic recombination generates either a pair of GFP^+^ *Trp53,Nf1*-null (TN-null) and sibling RFP^+^ WT OPCs, or a pair of yellow and colorless *Trp53,Nf1*-heterzygous OPCs, depending on the segregation pattern. In non-dividing cells (G0 or G1 phase of a cell cycle), NG2-Cre mediated recombination generates yellow *Trp53,Nf1*-heterozygous OPCs.

**Figure S2.**
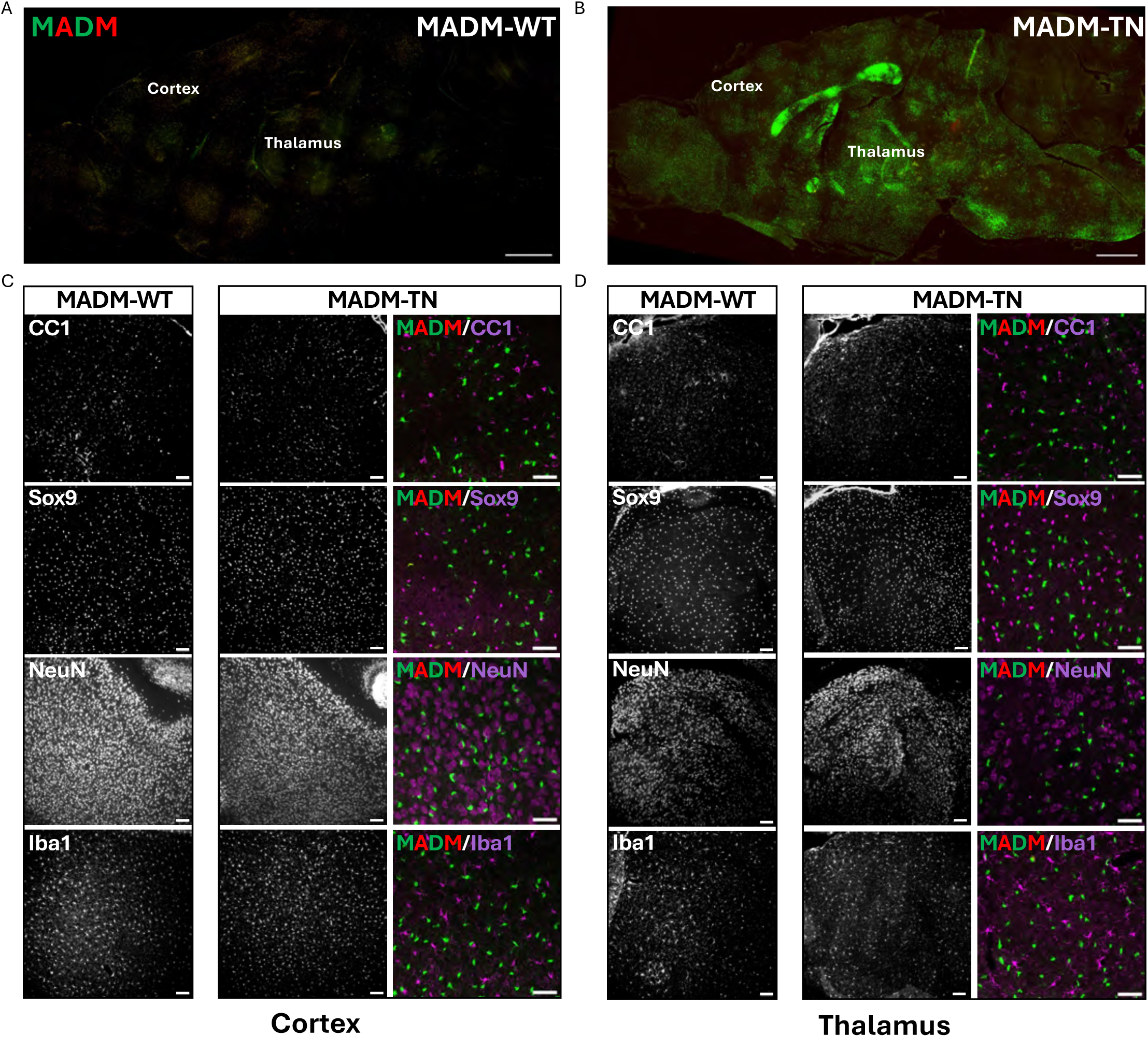
**TN-null OPCs do not grossly disrupt non-OPC brain resident cells. A-B**. Overview of MADM-WT (**A**) and MADM-TN (**B**) brains. Scale bar: 1000 μm. **C-D**. Distribution of oligodendrocytes (CC1^+^), astrocytes (Sox9^+^), neurons (NeuN^+^), microglia (Iba1^+^) was not disrupted in two representative brain regions – cortex (**C**) and thalamus (**D**) of MADM-TN mice. Scale bar: 100 μm. High-magnification images show that GFP^+^ TN-null OPCs intermingled with non-OPC brain resident cells (far right panel of **C, D**). Scale bar: 50 μm

**Figure S3.**
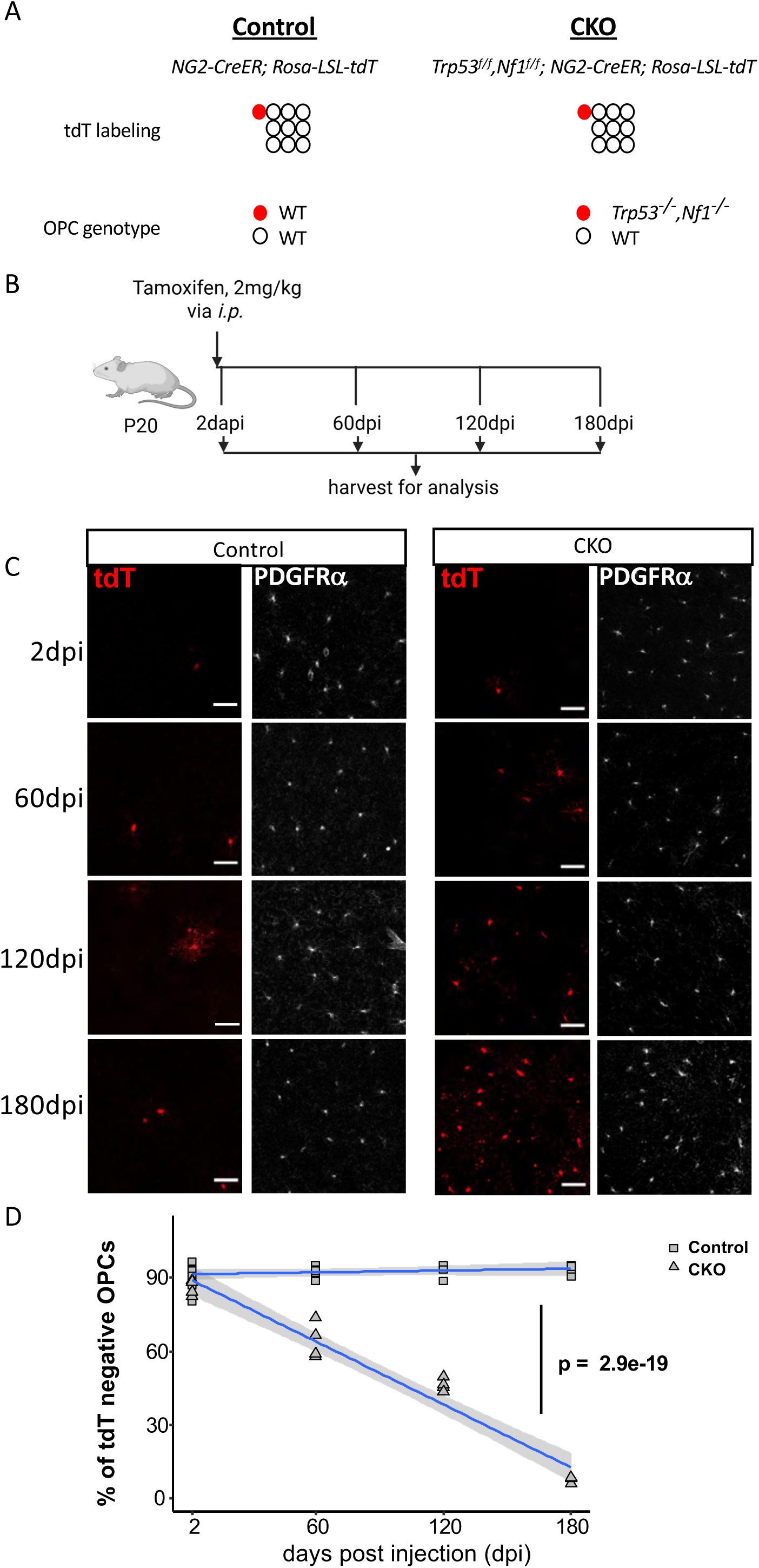
TN-null OPCs generated in adulthood also outcompete their neighboring normal OPCs. **A.** Genetic elements in the control and conditional knockout (CKO) mouse models, and the genotypes of tdT^+^ and tdT^-^ OPCs in each model. **B.** Tamoxifen regimen to generate sparsely labeled OPCs. Treated mice were harvested at 2, 60, 120 and 180dpi (days post injection) for analysis. **C.** Representative images of labeled OPCs in both control and CKO brains at each timepoint. Scale bar: 50 μm. **D.** Percentage of normal OPCs (tdT^-^) declines with post-injection day in the CKO but not the control mice. Temporal trends in OPC numbers from n ≥ 4 CKO and control animals per time point were modeled as a genotype-dependent exponential decay. The p value reports the statistical significance of the genotype term in the log-transformed linear model.

**Figure S4.**
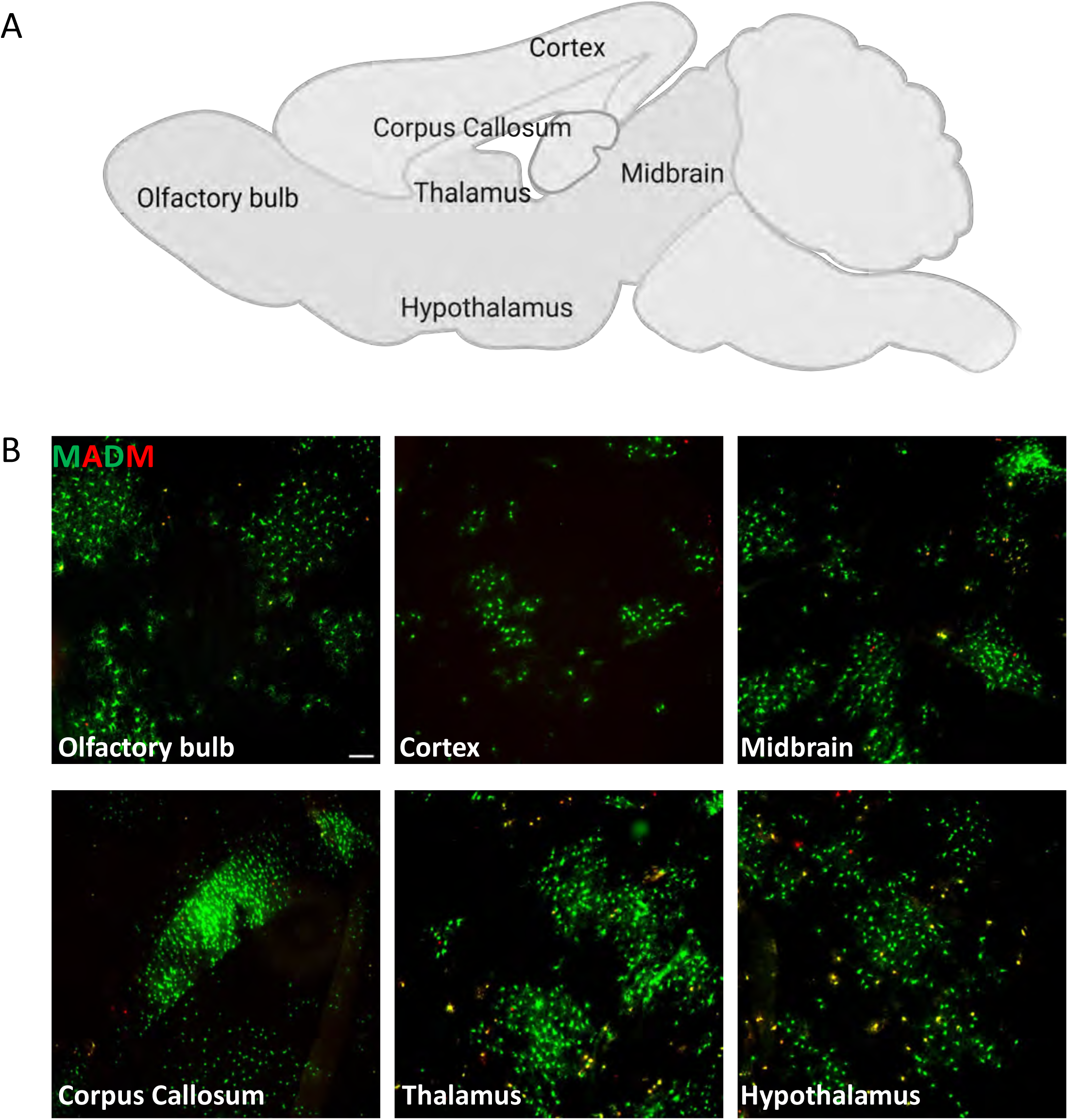
Clonal expansion of TN-null OPCs across all brain regions. **A.** Diagram of the brain regions examined in this study. **B.** Representative images of GFP^+^ TN-null OPC clones in various brain regions of P20 MADM-TN mice. Scale bar: 100 μm.

**Figure S5.**
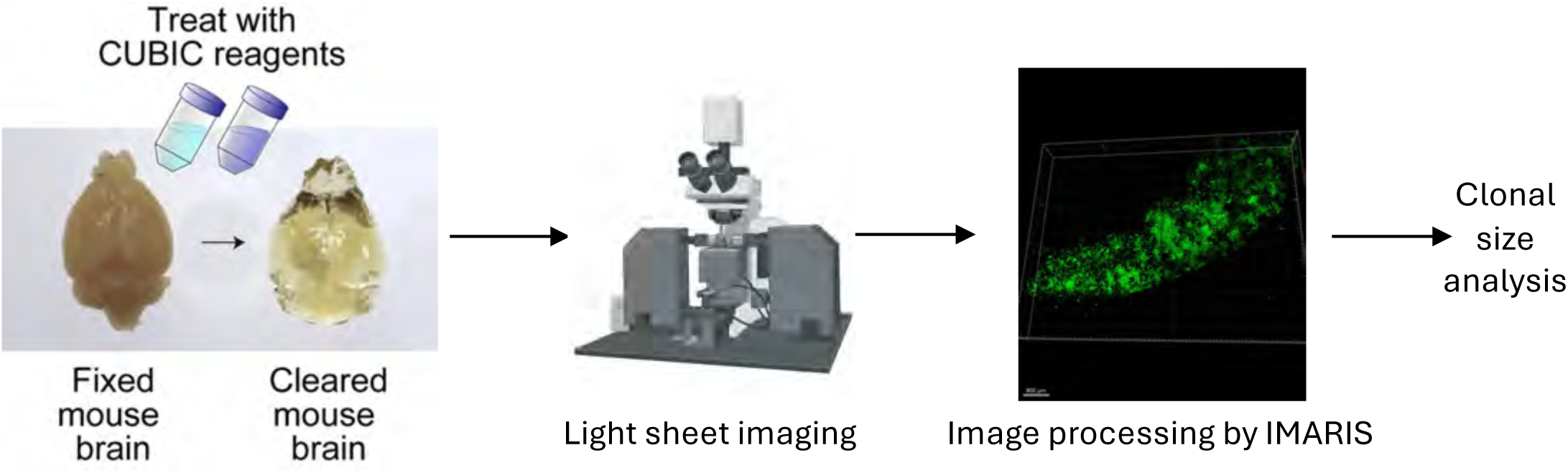
Experimental workflow for the 3-dimensional imaging of TN-null OPC clones in the brain.

**Figure S6.**
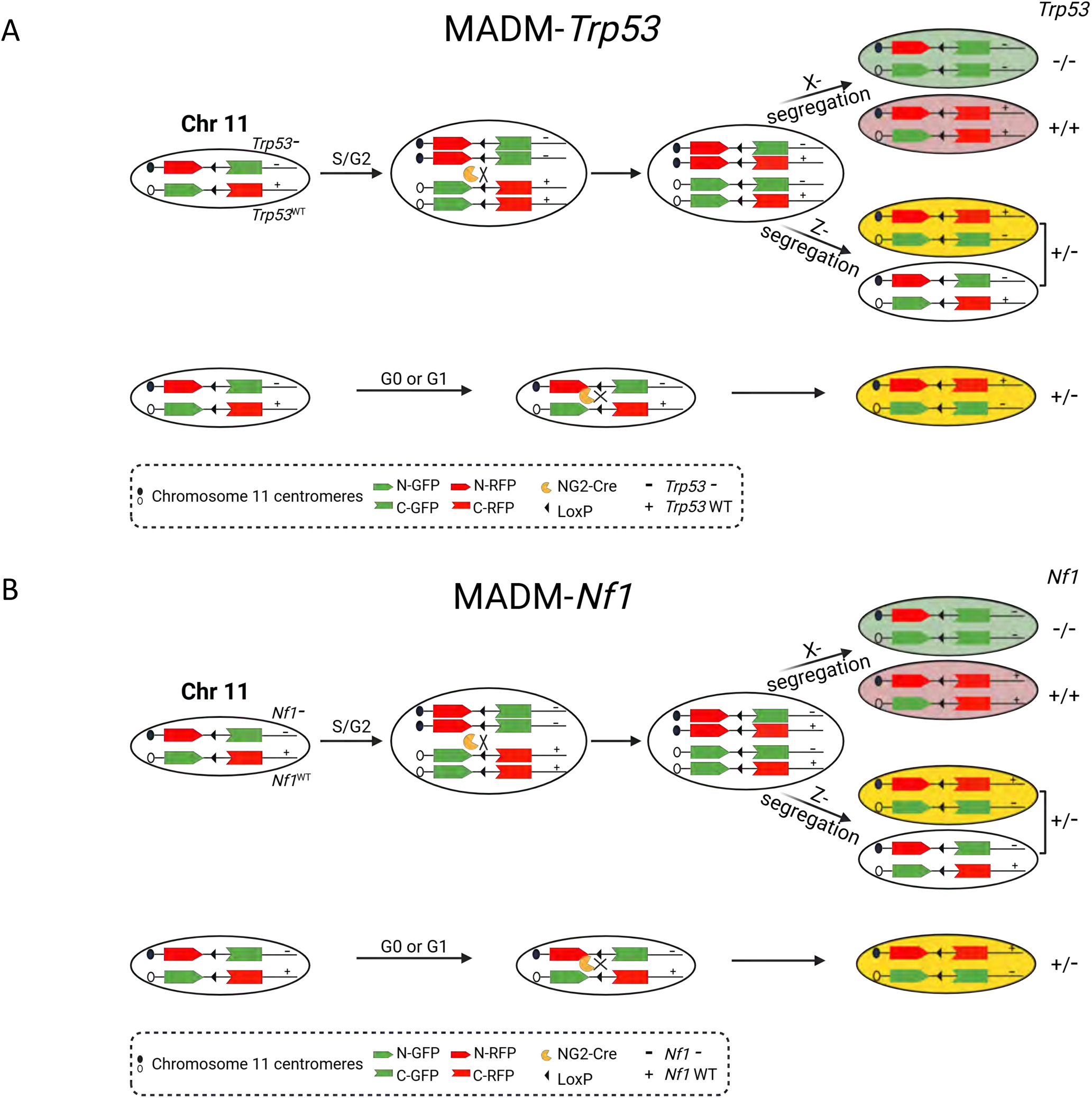
Detailed schematic of MADM model with individual mutations. **A.** Schematic of MADM-*Trp53* model. From an otherwise heterozygous, non-labeled mouse, NG2-Cre mediated mitotic recombination generates either a pair of GFP^+^ *Trp53*-null and sibling RFP^+^ WT OPCs, or a pair of yellow and colorless *Trp53*-heterzygous OPCs, depending on the segregation pattern. In non-dividing cells (G0 or G1 phase of a cell cycle), NG2-Cre mediated recombination generates yellow *Trp53*-heterzygous OPCs. **B.** Schematic of MADM-*Nf1* model. From an otherwise heterozygous, non-labeled mouse, NG2-Cre mediated mitotic recombination generates either a pair of GFP^+^ *Nf1*-null and sibling RFP^+^ WT OPCs, or a pair of yellow and colorless *Nf1*-heterzygous OPCs, depending on the segregation pattern. In non-dividing cells (G0 or G1 phase of a cell cycle), NG2-Cre mediated recombination generates yellow *Nf1*-heterzygous OPCs.

**Figure S7.**
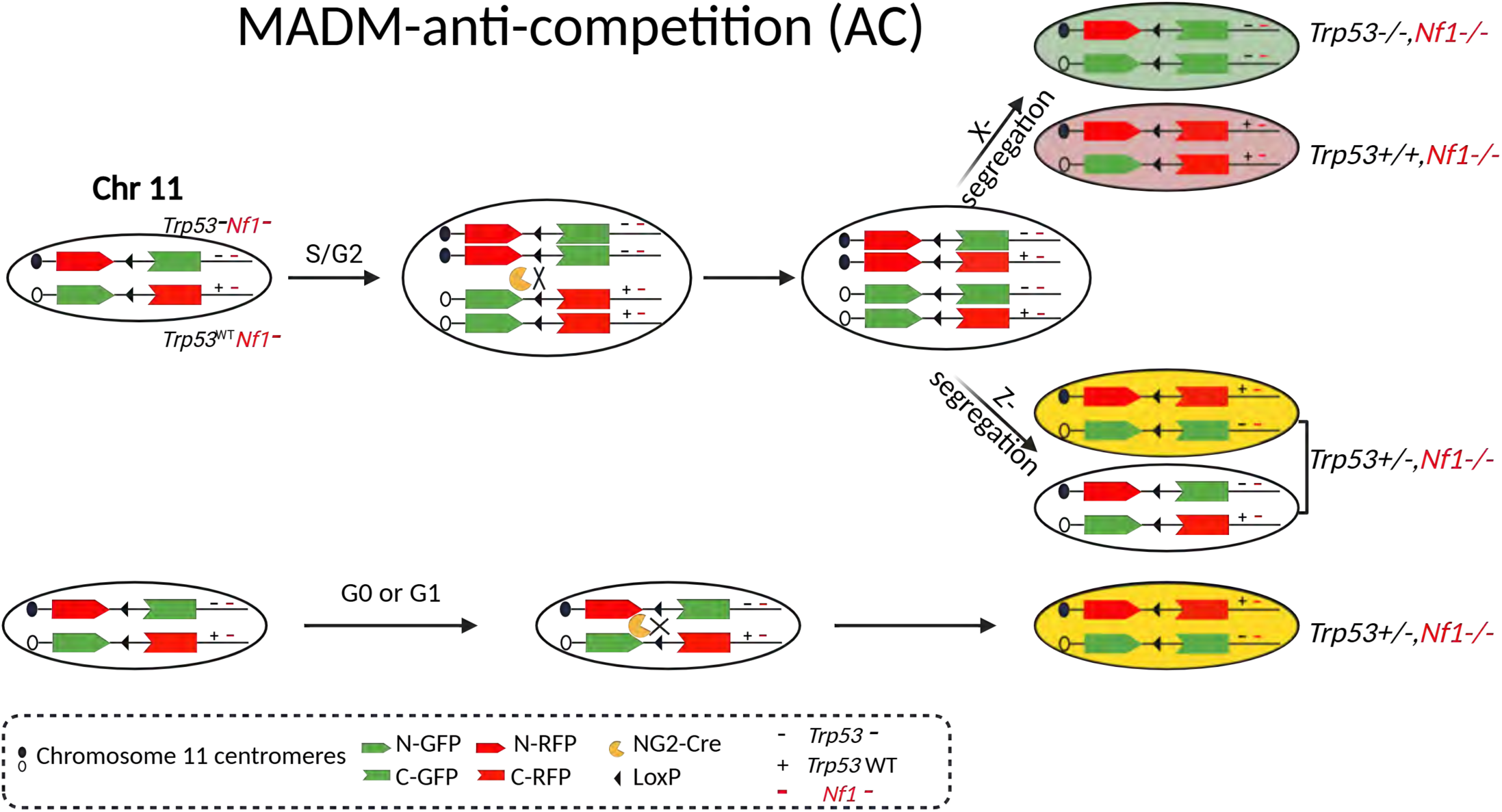
**Detailed schematic of generating MADM-anti-competition (AC)**. Schematic of MADM-anti-competition model, in which GFP^+^ OPCs are *Trp53,Nf1*-null while all other OPCs are *Nf1*-null.

**Figure S8.**
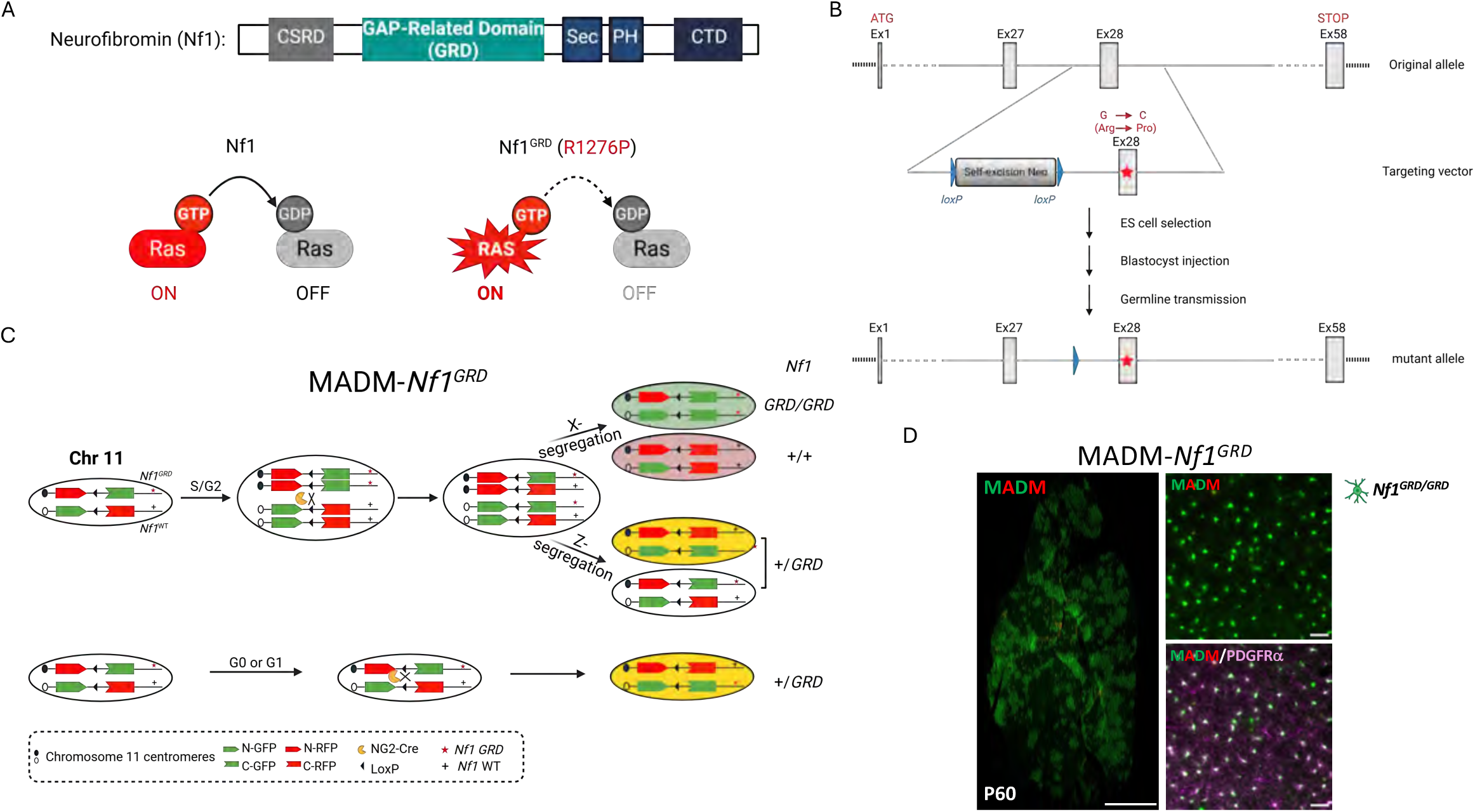
MADM-*Nf1^GRD^* model phenocopies the OPC competition of the MADM-*Nf1* model i. **A.** Schematic of protein domains in Nf1 including GAP-related domain (GRD) which acts as a negative regulator of Ras. Arg to Pro mutation at amino acid 1276 inactivates Nf1 GAP activity. **B.** Flowchart showing the process of generating a knock-in allele of *Nf1* (*Nf1^GRD^*) with a point mutation (G to C) resulting Arg to Pro mutation at amino acid 1276. **C.** Schematic of the MADM-*Nf1^GRD^* model. **D.** *Nf1-GRD*/*Nf1-GRD* OPCs outcompeted normal OPCs, phenocopying *Nf1*-null OPCs.

**Figure S9.**
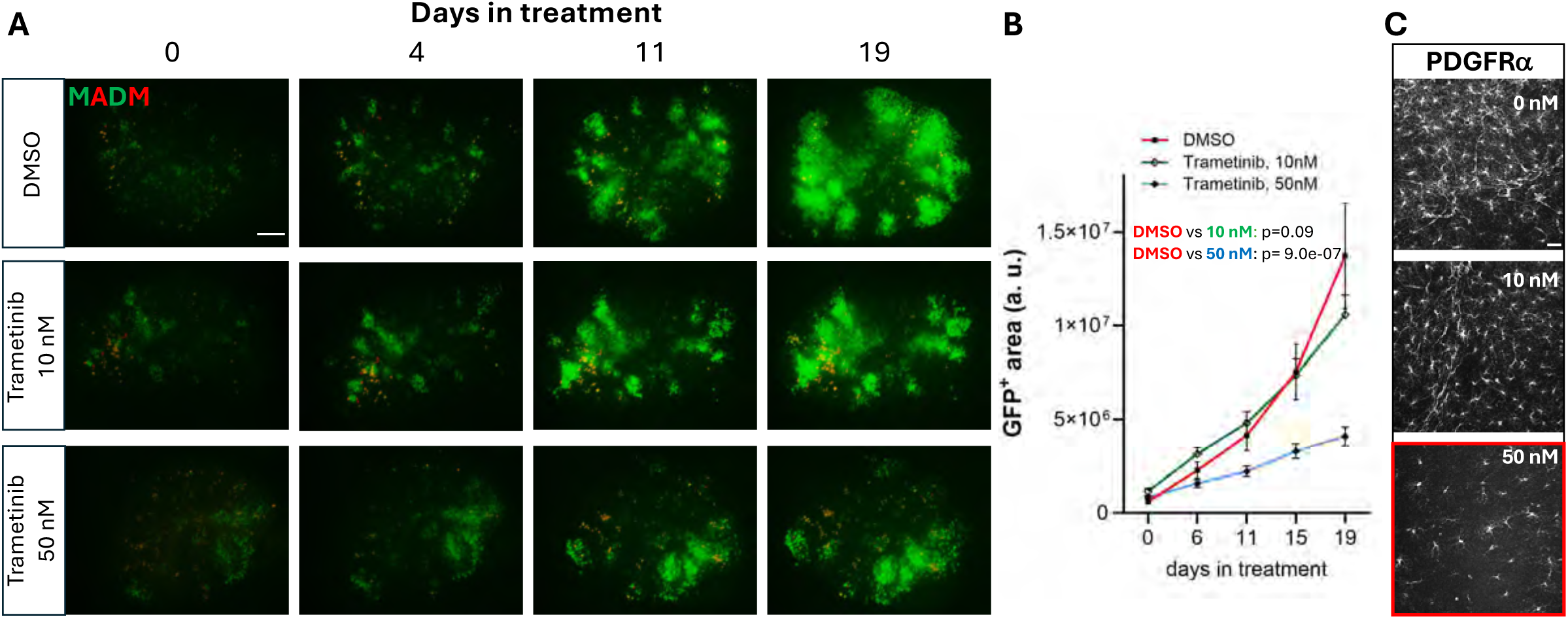
Trametinib, a MEK inhibitor, fails to block the expansion of TN-null OPCs without general toxicity to OPCs in *ex vivo* brain slice culture. **A.** GFP^+^ TN-null OPCs expanded in *ex vivo* cultured hippocampal slices (top). Addition of Trametinib at 10 nM failed to block the expansion of TN-null OPC (middle) while Trametinib at 50nM partially blocked the expansion of TN-null OPC (bottom). Scale bar: 200 µm. **B.** Line graph quantifying the expansion of GFP^+^ TN-null OPC along the days post-treatment. Temporal trends in GFP area from n ≥ 8 slides per group from MADM-TN animals per time point were modeled as a group-dependent exponential growth. The p value reports the statistical significance of the group term in the log-transformed linear model. Representative images of OPC density in brain slices at the time of harvest. 50 nM Trametinib treatment revealed dramatic loss of OPCs compared with DMSO or 10 nM Trametinib treatment, suggesting the lack of expansion is due to general toxicity rather than specific blockade of OPC competition. Scale bar: 50 μm.

**Figure S10.**
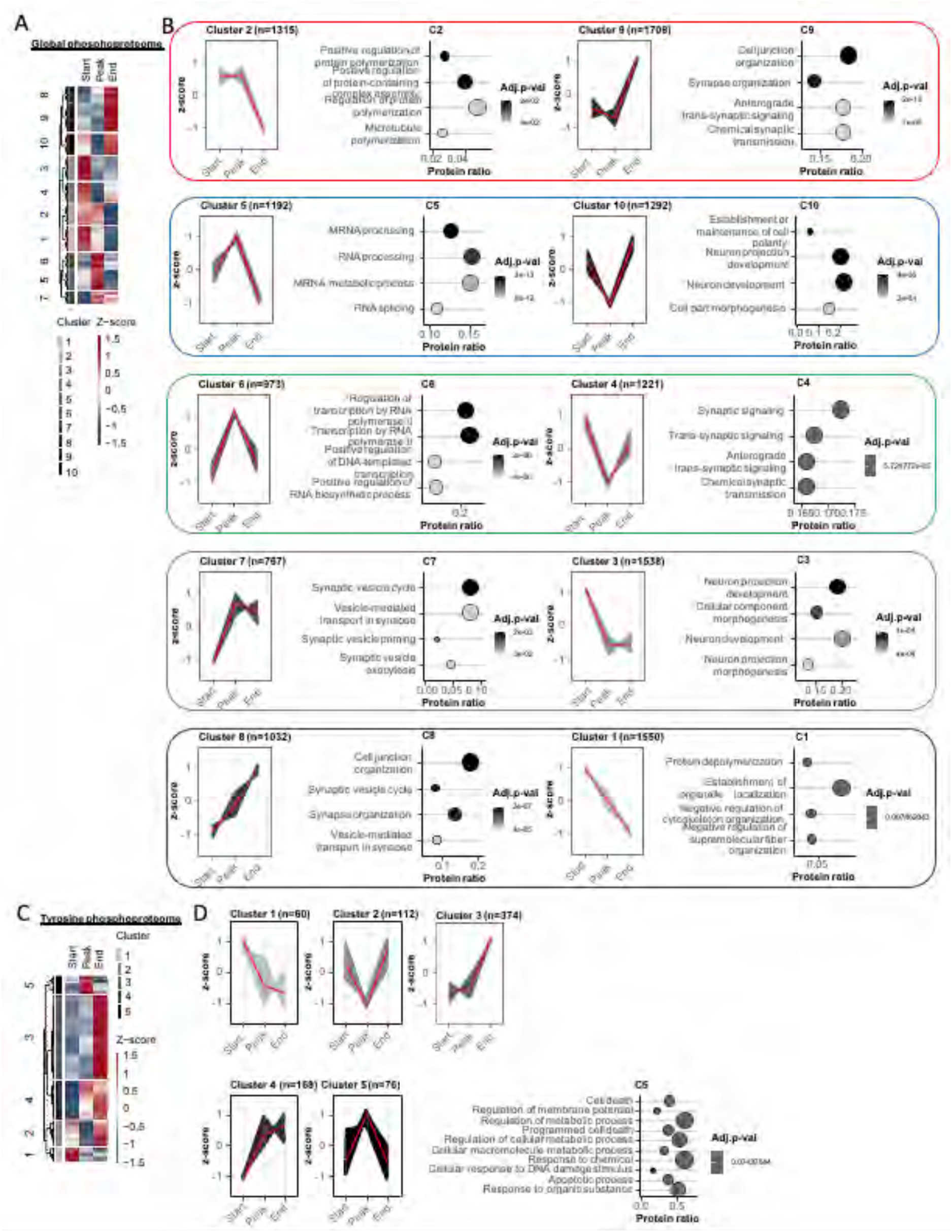
Analysis of normalized phosphoproteomic signals from MADM-TN brains at different stages of OPC competition. **A.** Phosphopeptides quantified by global phosphoproteomic analysis (mostly phosphoserine and phosphothreonine) were clustered using k-means clustering, leading to the identification of 10 distinct co-regulated clusters. **B.** The gene ontology term enrichment analysis of each cluster was performed using EnrichR. **C.** Phospho-tyrosine peptides were clustered using k-means clustering to identify 5 distinct co-regulated clusters. **D.** Cluster #5 had distinct enrichment of select gene ontology terms when analyzed by EnrichR.

**Figure S11.**
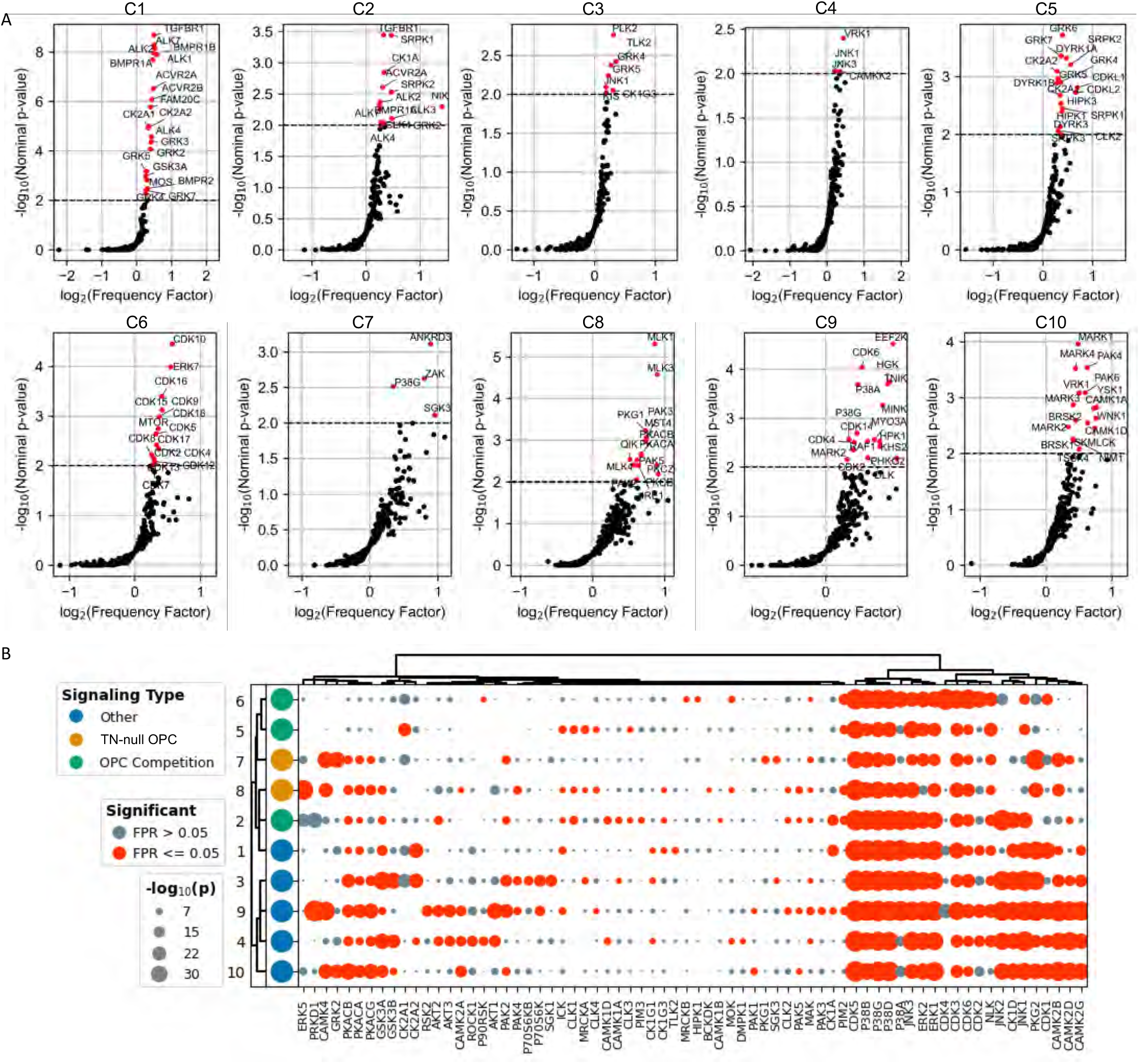
Prediction of active kinases in each cluster. **A.** Kinase Library and **B.** KSTAR analysis were used to predict kinases responsible for phosphopeptides in each cluster in Figure S10**A-B**.

**Figure S12.**
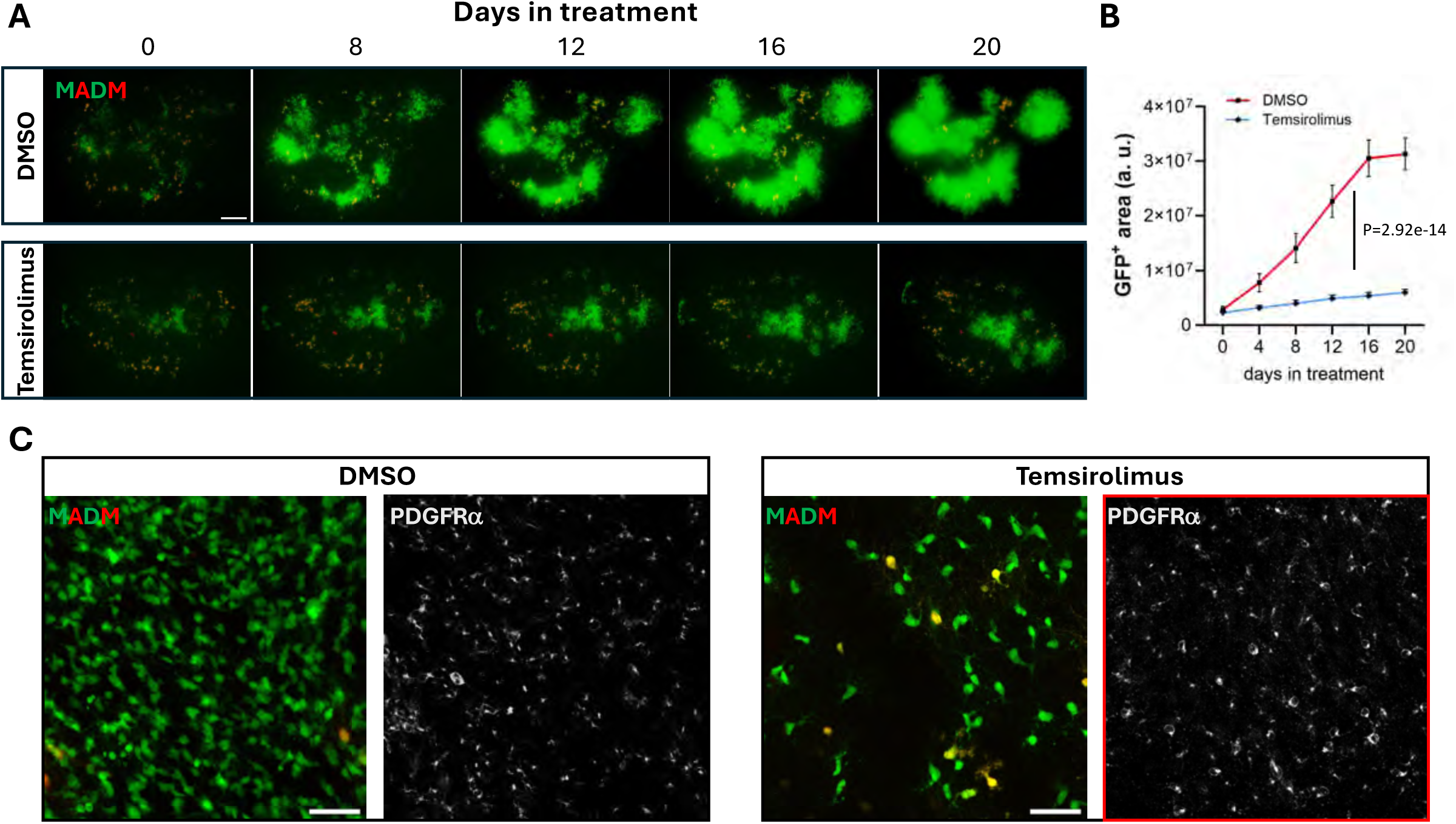
Temsirolimus, a mTORC1 inhibitor, specifically blocks the expansion of TN-null OPCs without general toxicity to OPCs in *ex vivo* brain slice culture. **A.** GFP^+^ TN-null OPCs expanded in *ex vivo* cultured hippocampal slices (top). Treatment with Temsirolimus, a mTORC1 inhibitor, blocked the expansion of TN-null OPCs (bottom). Scale bar: 200 µm. **B.** Line graph quantifying the expansion of GFP^+^ TN-null OPC along the days post-treatment. Temporal trends in GFP area from n ≥ 8 slides per group from MADM-TN animals per time point were modeled as a group-dependent exponential growth. The p value reports the statistical significance of the group term in the log-transformed linear model. **C.** Representative images of GFP^+^ TN-null OPCs (left) and total OPCs (right) in control and Temsirolimus-treated brain slices at the time of harvest. Scale bar: 50 μm.

**Figure S13.**
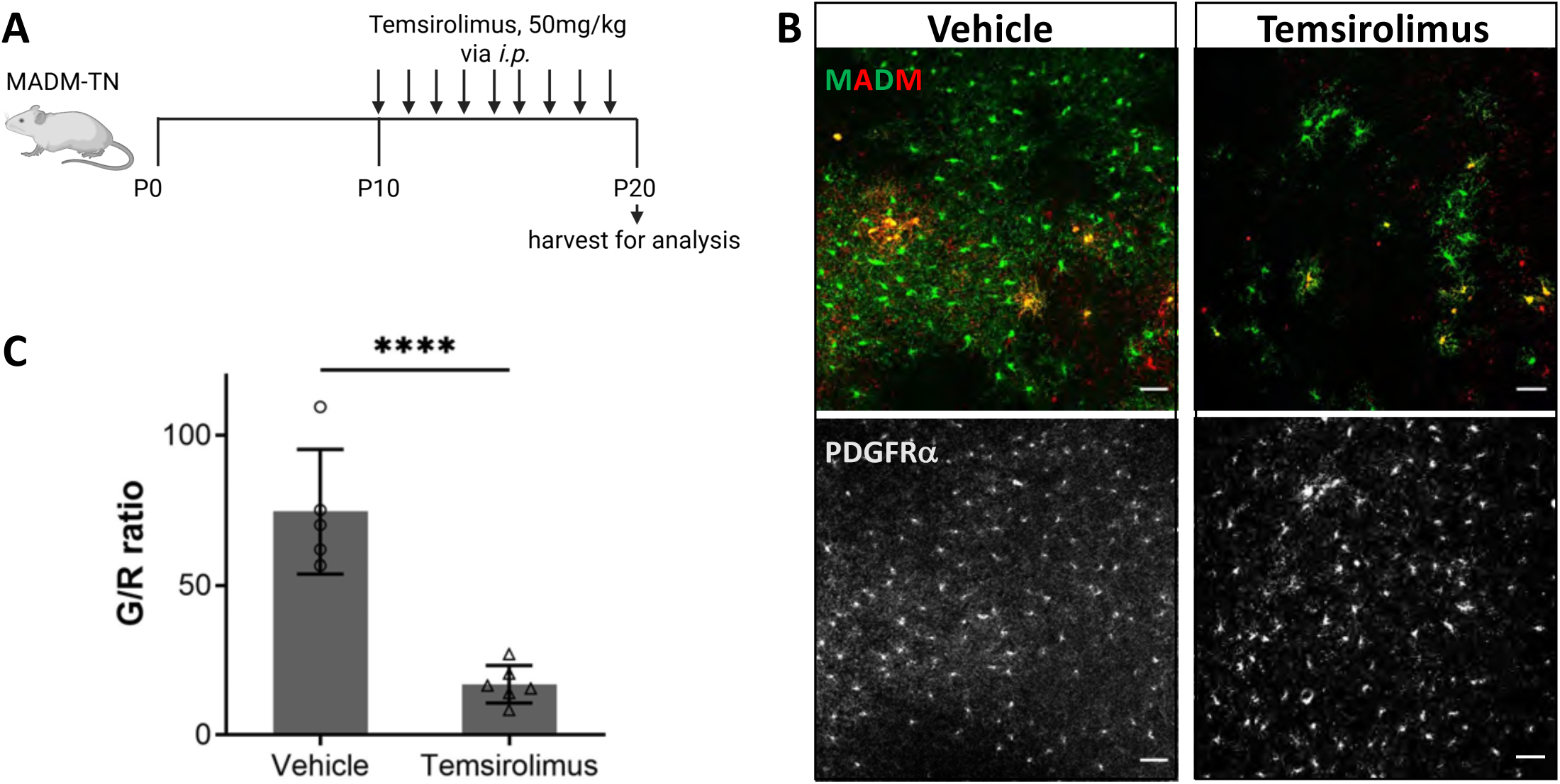
Temsirolimus blocks OPC competition in the MADM-TN model. **A.** Temsirolimus treatment regimen of MADM-TN mice. **B.** Representative images of GFP^+^ TN-null OPCs (top) and total OPCs (bottom) in both vehicle and Temsirolimus-treated brains. Scale bar: 50 μm. **C.** Bar graph showing the G/R ratio at the end of treatment. Pairwise ratio t-test was performed, ****p<0.0001. ≥5 MADM-TN brains from each group were used for quantification.

**Figure S14.**
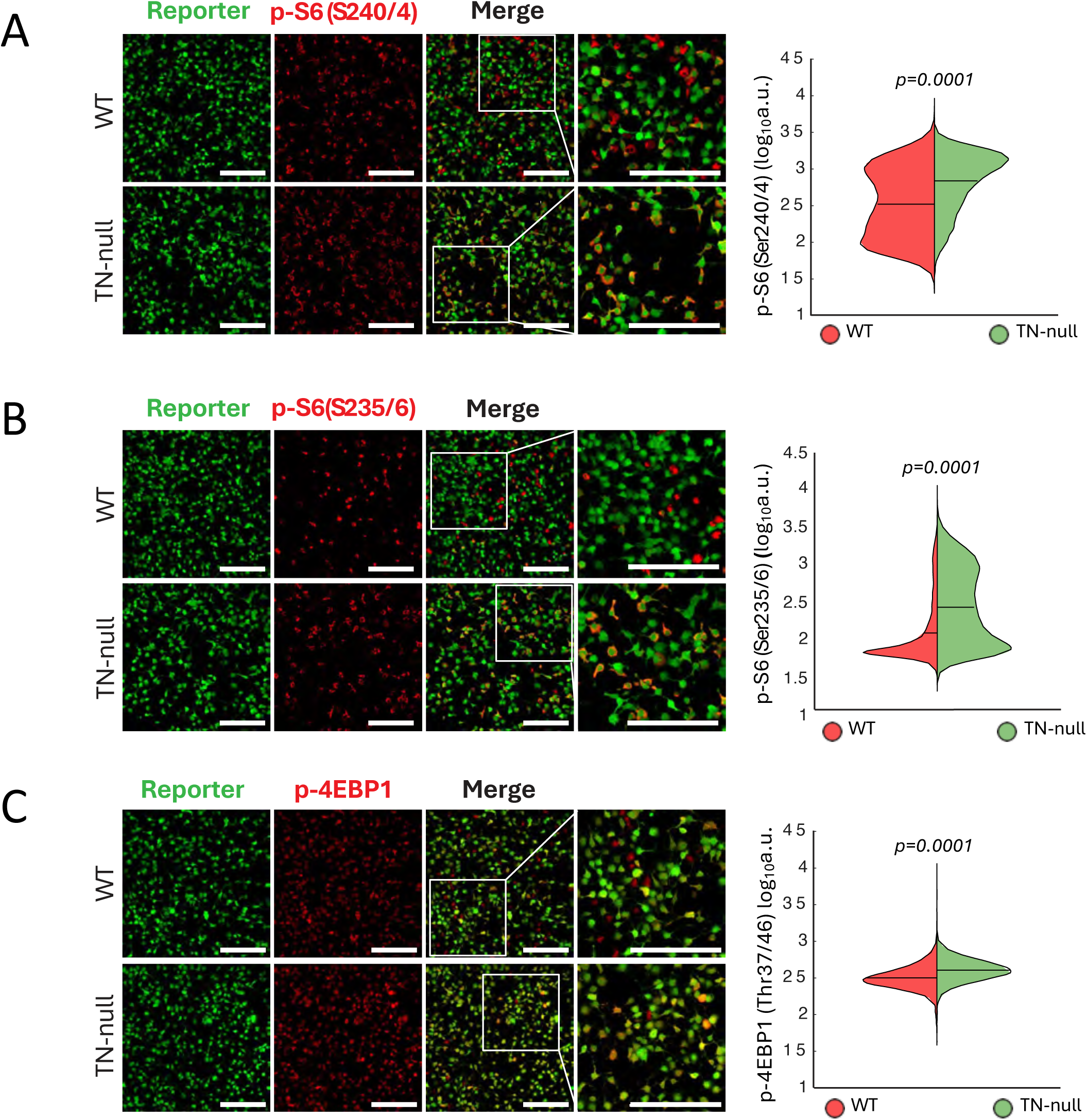
Single cell analysis of mTORC1 activity in acutely purified DKO and WT OPCs. Representative immunofluorescence images on the left showing the OPCs (with reporter) and the staining of mTORC1 substrates: p-S6 S240/4 (**A**) p-S6 S235/6 (**B**), and p-4EBP1 T37/41 (**C**) respectively. Scale bars: 200 µm. Violin plots on the right summarizing the quantitative analysis of signal intensity for each mTORC1 substrate at the single cell level (from total of 10,000 cells pooled from replicates without bias). Statistical significance was determined by a one-sided permutation test.

**Figure S15.**
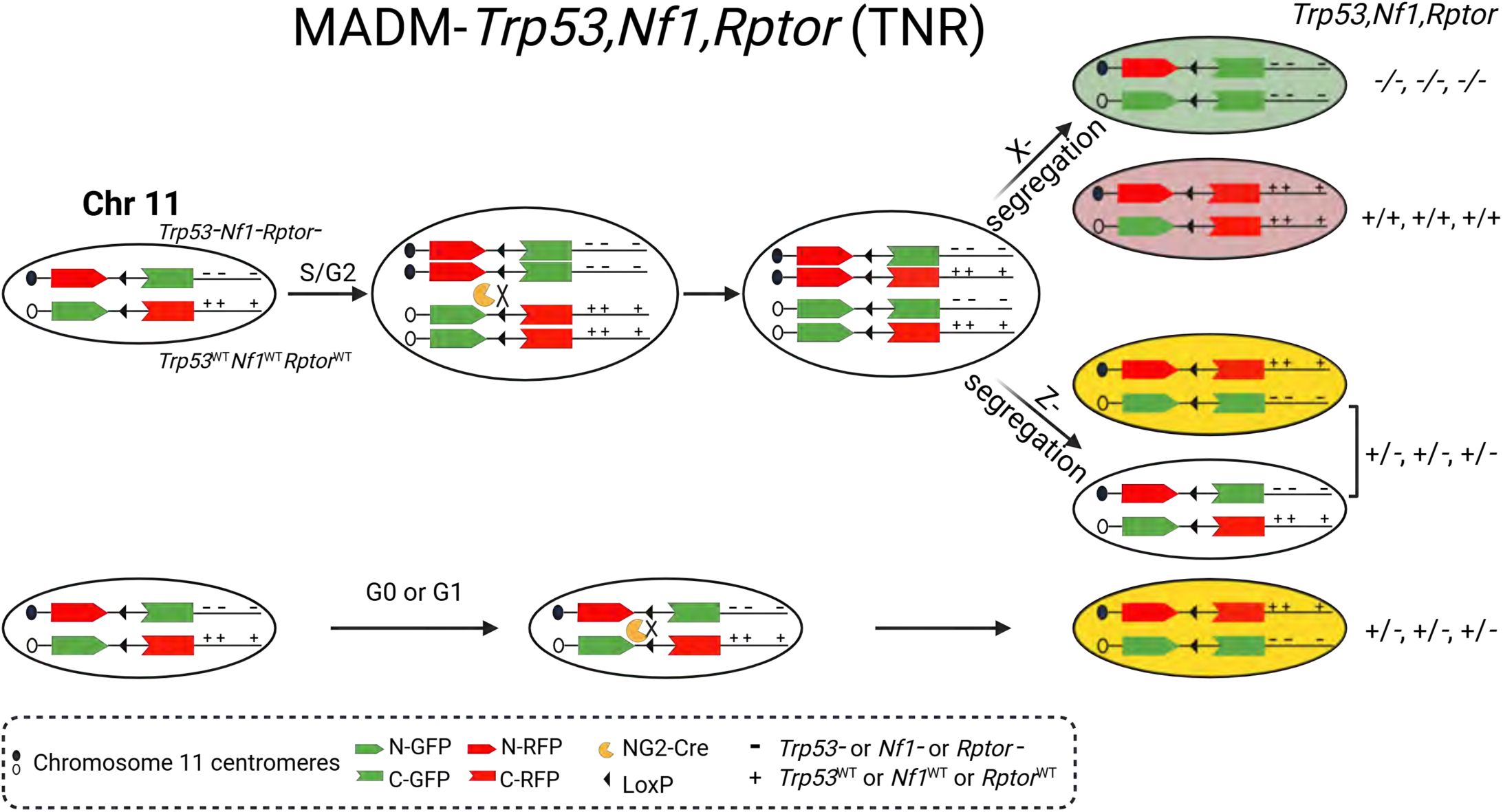
Detailed schematic of MADM-*Trp53,Nf1,Rptor* (TNR) model. From an otherwise heterozygous, non-labeled mouse, NG2-Cre mediated mitotic recombination generates either a pair of GFP^+^ *Trp53,Nf1,Rptor*-null (TNR-null) and sibling RFP^+^ WT OPCs, or a pair of yellow and colorless *Trp53,Nf1*-heterzygous OPCs, depending on the segregation pattern. In non-dividing cells (G0 or G1 phase of a cell cycle), NG2-Cre mediated recombination generates yellow *Trp53,Nf1,Rptor*-heterzygous OPCs.

**Figure S16.**
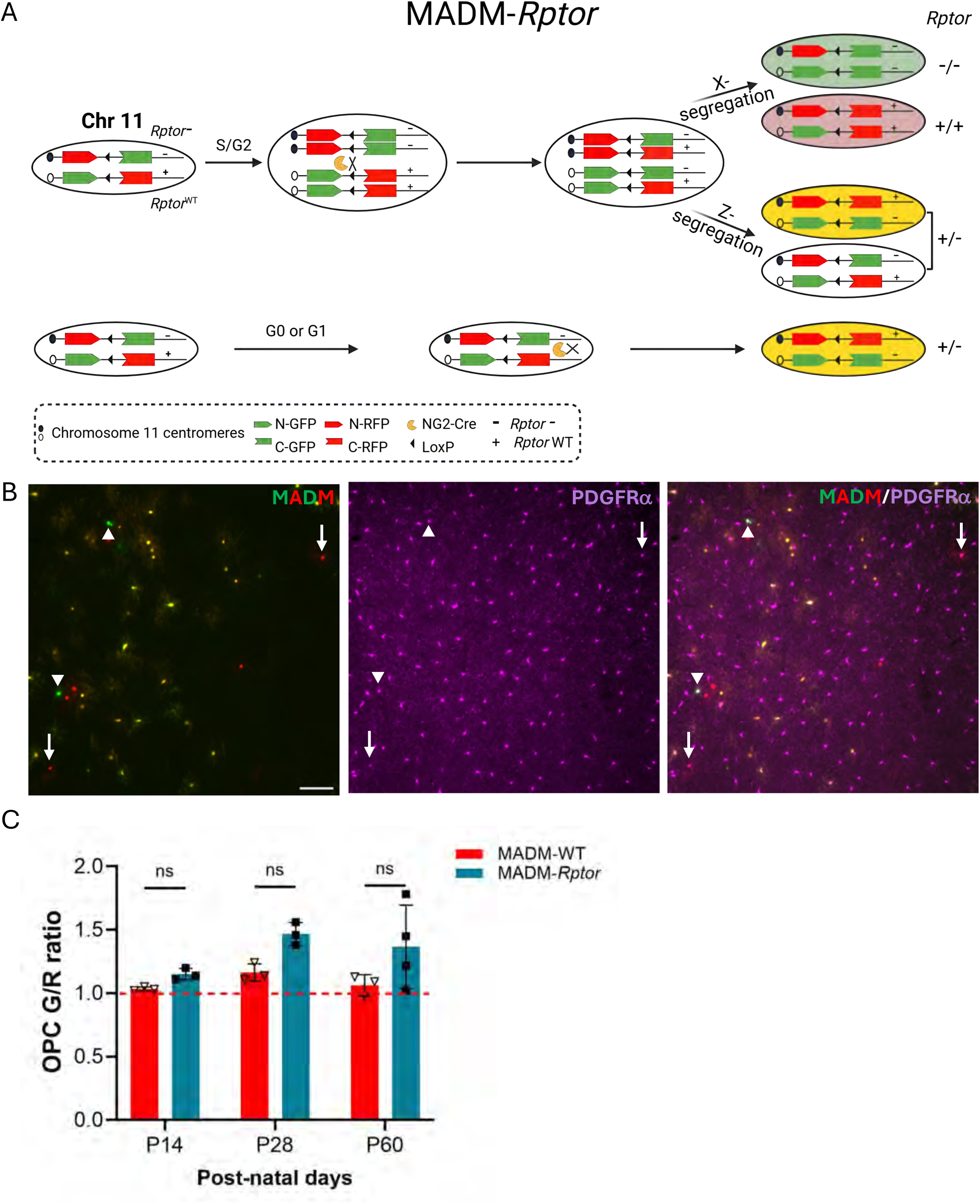
mTORC1 is dispensable for OPC development and maintenance. **A.** Schematic of MADM-*Rptor* model. **B.** High-magnification images to identify GFP^+^ *Rptor*-null OPCs (white arrow-heads) and RFP^+^ *Rptor*-WT OPCs (white arrows) for G/R ratio quantification. Scale bar: 50 μm. **C.** Bar graph showing G/R ratio remained ∼1 at various ages. Pairwise ratio t-test was performed, ns p>0.05. ≥3 MADM-*Rptor* and MADM-WT brains from each age group were used for quantification.

**Figure S17.**
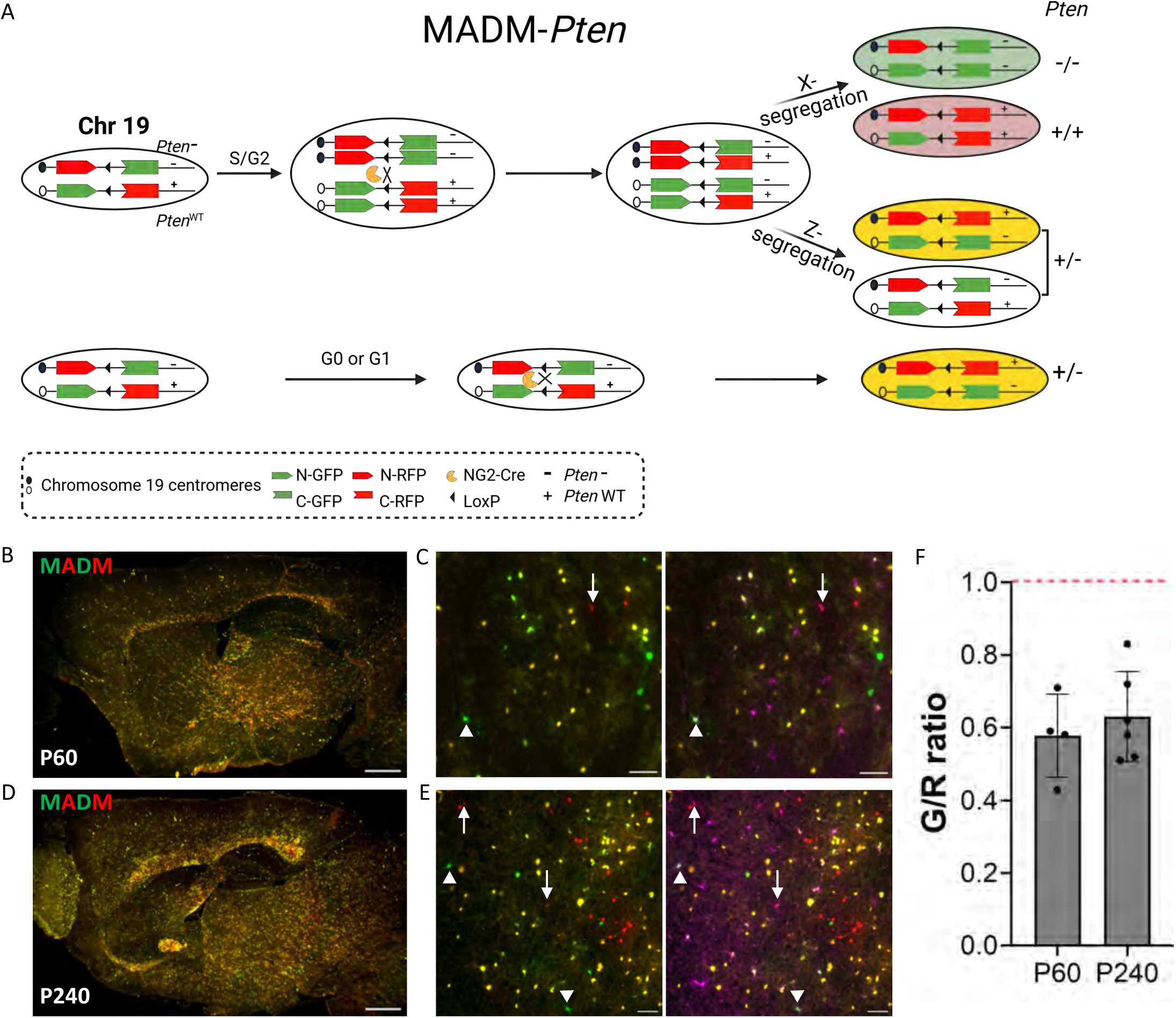
*Pten* loss is not sufficient to drive OPC competition. **A.** Schematic of the MADM-*Pten* model. **B-E**. Macroscopic images showing the lack of expansion of GFP^+^ *Pten*-null OPCs at both P60 (**B**) and P240 (**D**). Scale bar: 500 μm. **C,E** high-magnification images to identify GFP^+^ *Pten*-null OPCs (white arrow-heads) and RFP^+^ *Pten*-WT OPCs (white arrows) for G/R ratio quantification. Scale bar: 50 μm. **F.** Bar graph showing G/R ratio lower than 1 at both P60 and P240. ≥4 MADM-*Pten* brains from each age group were used for quantification.

**Figure S18.**
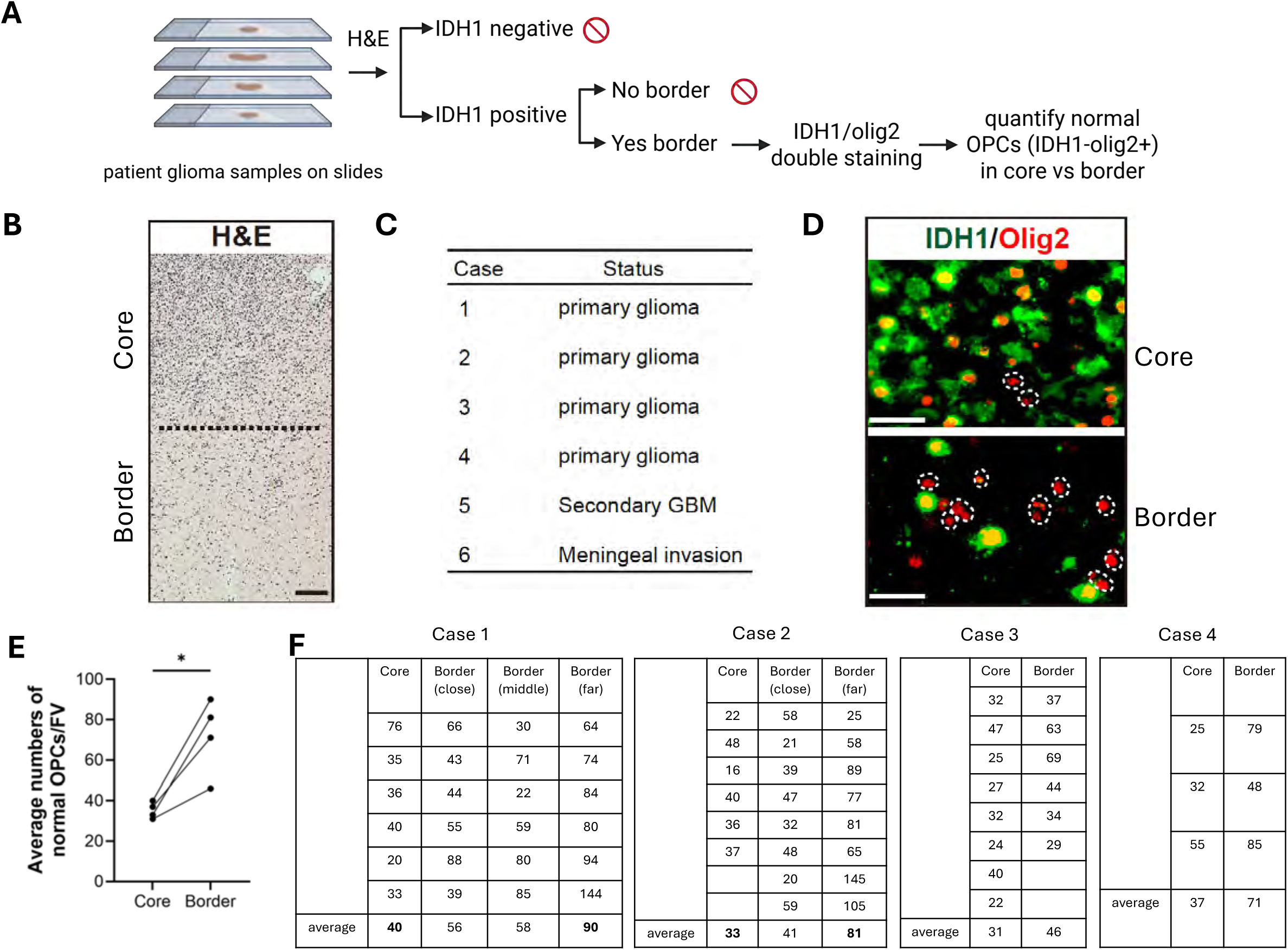
Evidence of OPC competition in IDH-mutant patient samples. **A.** Workflow of identifying are IDH-mutant glioma patient samples that had both tumor core and border regions for further analysis. **B.** Representative image showing IDH-mutant glioma that contains both tumor core and border regions, based on H&E staining. Scale bar: 100 μm. **C.** Six IDH-mutant glioma samples that met the selection criteria for this experiment were picked from 111 cases. **D.** Representative images showing the staining patterns of IDH1 and olig2 at tumor core and border regions, and the quantification of normal OPCs in each region (dotted circles: olig2^+^IDH1^-^). Scale bar: 30 μm. **E.** Pair-wise analysis of the average number of normal OPCs per field of view (FV) showed that four cases had reduced number of normal OPCs in tumor core compared to border regions. In cases that have multiple sampling in border regions, the furthest border was used for plotting. Considering the tumor heterogeneity, we quantified 3-8 FVs depending on the size of core/border and used the average value for plotting. *p<0.05, paired t-test was performed. **F.** Raw data of the normal OPC quantification in all 4 cases in (**E**). The average number of normal OPCs in tumor cores were similar. For cases with multiple borders, we observed that the further away from tumor core, the larger the number of normal OPCs.

## Supplement video

GFP^+^ TN-null OPC clones from the cortical region of P21 MADM-TN brain were shown here.

